# The binding mechanism of *Streptococcus suis* accessory virulence factor and adhesin SadP to globotetraosylceramide

**DOI:** 10.1101/2020.05.29.122762

**Authors:** Miralda Madar Johansson, Eva Bélurier, Anastassios C. Papageorgiou, Anders P. Sundin, Jani Rahkila, Teemu Kallonen, Ulf J. Nilsson, Santeri Maatsola, Thomas K.M. Nyholm, Jarmo Käpylä, Jukka Corander, Reko Leino, Jukka Finne, Susann Teneberg, Sauli Haataja

## Abstract

*Streptococcus suis* is part of the pig commensal microbiome and a major pathogen causing pneumonia and meningitis in pigs and occasionally also zoonotic meningitis. According to genomic analysis, *S. suis* is divided into asymptomatic carriage, respiratory and systemic strains with distinct genomic signatures. The virulence factor *S. suis* adhesin P (SadP) recognizes the galabiose Galα1–4Gal-oligosaccharide. Based on its oligosaccharide fine specificity, SadP can be divided into subtypes P_N_ and P_O_. We show here that subtype P_N_ is distributed in the systemic strains that cause meningitis, whereas type P_O_ is found in asymptomatic carriage and respiratory strains. Both types of SadP are shown to predominantly bind to pig lung globotriaosylceramide (Gb3). However, SadP adhesin from systemic subtype P_N_ strain also binds to globotetraosylceramide (Gb4). Mutagenesis studies of the galabiose-binding domain of type P_N_ SadP adhesin showed that the amino acid asparagine-285, which is replaced by an aspartate residue in type Po SadP, was required for binding to Gb4 and, strikingly, it was also required for interaction with the glycomimetic inhibitor phenylurea-galabiose. Molecular dynamics simulations provided further insight into the role of Asn-285 for Gb4 and phenylurea-galabiose binding, suggesting additional hydrogen bonding to terminal GalNAc of Gb4 and urea-group. Thus, the Asn-285-mediated molecular mechanism of type P_N_ SadP binding to Gb4 could be used as a candidate to selectively target *S. suis* in invasive systemic disease without interfering with commensal strains, which may open up new venues for developing intervention strategies against this pathogen.

## Introduction

Bacterial adhesion to host cell surfaces is a prerequisite for infectious disease. Host cell surfaces are heavily covered by surface carbohydrates that form the glycocalyx layer (1, 2). Pathogenic bacteria and their toxins can treacherously exploit carbohydrates to attach and invade cells. One example is a group of endogenous glycosphingolipids, known as globo series membrane glycosphingolipids, which are grouped based on the Galα1–4Gal (galabiose) structure. Globo series glycolipids are cellular receptors for toxic action of Verotoxins and for the attachment of bacteria, such as uropathogenic *Escherichia coli*, *Streptococcus suis* and *Pseudomonas aeruginosa* (3–6).

Since antibiotics are increasingly losing their power, alternative antimicrobial compounds need to be invented and developed (7, 8). Transformation of bacteria to antibiotic-resistant forms can be overcome by developing compounds that block *in vivo* colonization and virulence factors without killing the pathogens. Cell surface carbohydrate glycomimetics are the most tempting pipeline for the development of new generation of antimicrobials (9, 10).

*Streptococcus suis* is found as a commensal bacterium in all pigs and it is an important pig pathogen causing septicemia, pneumonia and meningitis. It also causes severe zoonotic meningitis. Based on clinical outcome, the *S. suis* strains can be classified as asymptomatic carriage, respiratory and systemic strains. Recent data show that *S. suis* produces an alarmingly vast array of factors causing high antibiotic resistance (11). Genomic analysis shows that the asymptomatic carriage and systemic strains have clear differences in their core and accessory genomes, whereas respiratory strains have more overlapping features (12). This could suggest that environmental factors such as overcrowding and synergistic activities with other keystone pathogens (13), can alter pigs to develop severe symptoms and pneumonia. Instead, the virulence genes are enriched in systemic strains and their accessory genomes contribute to the invasive *S. suis* infections and meningitis (12). There is multiple and redundant colonization and virulence factors required for *S. suis* disease. The distribution of virulence factors is also different depending on the geographical regions impacting the accessory genomes, which hampers the development of effective vaccines (14). In order to develop new therapeutics and vaccines the molecular mechanisms of accessory virulence factors need to be resolved in atomic level.

*S. suis* strains expressing SadP adhesin bind specifically to Galα1–4Gal-containing oligosaccharides (15). Interestingly, galabiose-binding strains can be divided to subtypes P_N_ and P_O_ based on how they recognize galabiose-containing carbohydrates in hemagglutination assays (16). Type P_N_ strains recognize galactose, *N*-Acetylgalactosamine, and the ceramide-linked oligosaccharides Gb3 (Galα1–4Galβ1–4Glc) and Gb4 (GalNAcβ1–3Galα1–4Galβ1–4Glc). Subtype P_O_ recognizes only galactose and preferentially binds to Gb3. Recently, *S. suis* galabiose-binding adhesin SadP was identified as an LPNTG-anchored cell-wall protein (6). The carbohydrate interaction of SadP has been shown to mediate *S. suis* binding to pharyngeal epithelium and intestinal epithelial cells (16, 17). In an *S. suis* murine infection model, the mice deficient in globotriaosyl ceramide (Gb3) expression (knockout in alpha-1,4-galactosyltransferase A4GALT) developed less severe brain inflammation and injury (18). Recently, low picomolar concentrations of glycomimetic 3-phenylurea-galabiose-containing dendrimers were shown to block *S. suis* SadP (type P_N_) adhesin binding activity (19). However, the molecular basis of differences in galabiose-binding mechanisms of SadP adhesins have remained elusive. In the present study, we have characterized the distribution of SadP subtypes in *S. suis* strains and the molecular binding mechanism of type P_N_ and P_O_ SadP to globosyl oligosaccharides. The results show that *S. suis* SadP type P_N_ is distributed in the systemic strains and its specific N285-mediated binding mechanisms to globotetraosyl ceramide distinguishes it from type P_O_ sadP.

## Results

### Sequence analysis of SadP subtypes P_N_ and P_O_, structural modelling and distribution in *S. suis* systemic, respiratory and non-clinical strains

SadP adhesins are 80 kDa LPNTG-anchored cell-wall proteins, which contain the signal sequence for secretion, the N-terminal galabiose binding domain, tandem repeat domains and C-terminal LPNTG cell-wall anchor domain (6). The galabiose-binding domains have no significant sequence homologs to other bacterial proteins or to other carbohydrate binding domains. The SadP N-terminal galabiose-binding domains (aa125-328) of hemagglutinating type P_N_ and type P_O_ strains were cloned, sequenced and compared with multiple alignment (Fig 1A). The structure consists of three α-helixes and ten β-strands (β1-β10) that form a β-sandwich core domain. The sequences of the N-terminal galabiose-binding domain of type P_N_ SadP adhesins are 100 % identical and are found in virulent serotype 2 strains. There is more sequence variation in the galabiose-binding site of type P_O_ SadP. The carbohydrate binding sites of types P_N_ and P_O_ are highly conserved (Fig 1B). Interestingly there is a conservative change at position 285 from asparagine to aspartate, which could have an effect for the interaction of type P_N_ SadP with GalNAcβ1–3Galα1–4Gal-oligosaccharide.

**Fig 1.**
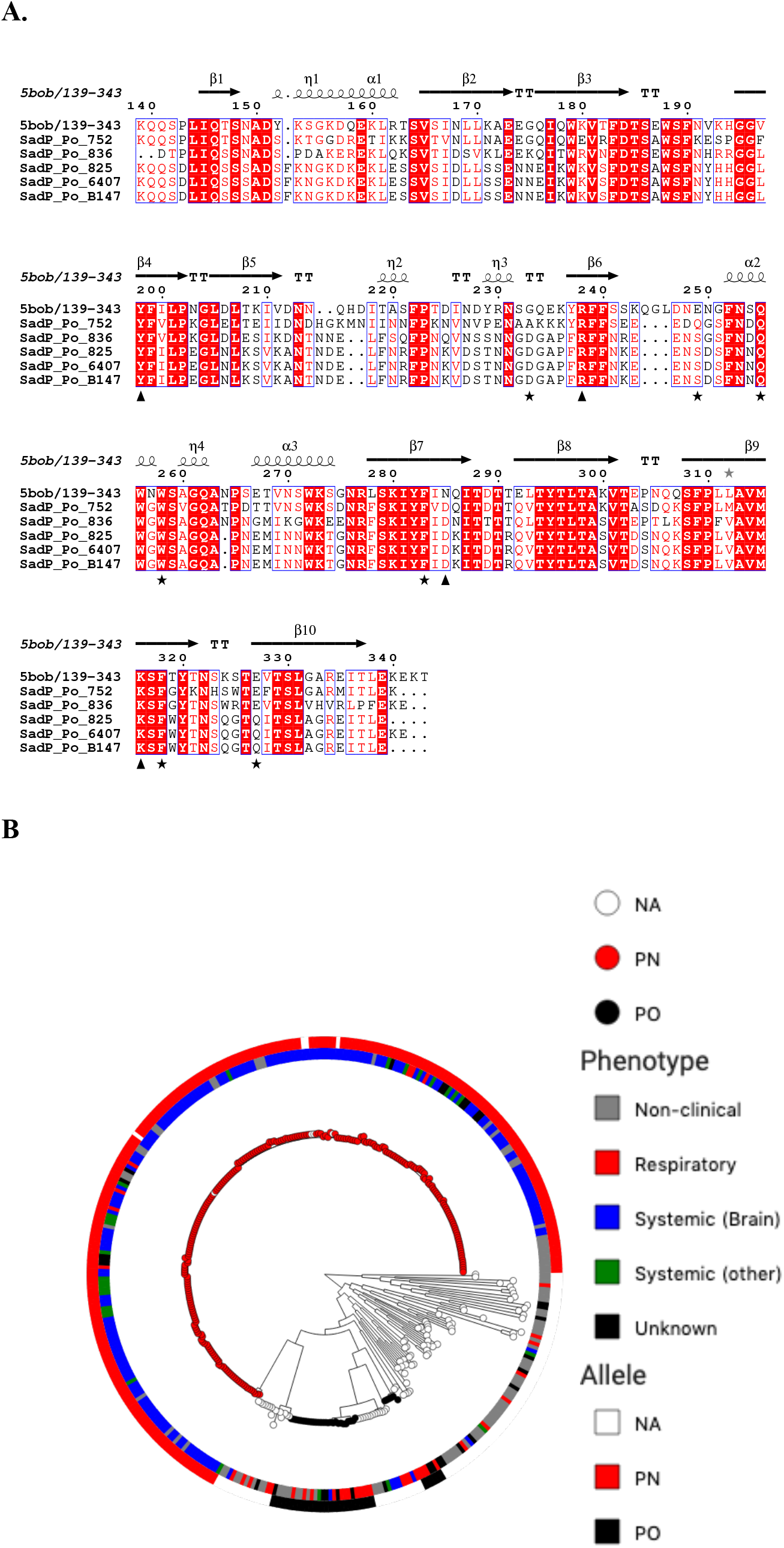
Homology of SadP types P_N_ and P_O_ galabiose-binding domains and their distribution in systemic, respiratory and non-clinical *S. suis* isolates. A. Multiple alignment of SadP galabiose binding domains of type P_N_ and type P_O_ strains. Conserved amino acids are shown in shaded red rectangles and white letters. Residues with >70% similarity according to their physicochemical properties are framed in white background and shown with red letters. Residues involved in hydrogen-bond formation with galabiose (Galα1–4Gal) are shown with (▴) and those involved in hydrophobic interactions with a star (★). Residues 139-343 of SadP are included as found in the crystal structure (pdb id 5BOB). Secondary structure elements are shown on the top. Figure was created with ESpript (46). **B. Distribution of SadP type P_N_ and P_O_ alleles in** *S. suis* **clinical and non-clinical isolates.** Maximum-likelihood core genome phylogeny of 374 *S. suis* isolates from Weinert et al. 2015. The phylogeny was run with RAxML using SNP sites. The nucleotide substitution model used was GTR GAMMA. Tips of the tree are coloured according to the SadP allele detected with SRST2. NA means that with the default parameters neither allele was detected. The inner circle shows the phenotype according to Weinert et al. 2015. Grey non-clinical, red respiratory, blue systemic (brain), green systemic (other) and black unknown. The outer circle is coloured according to the SadP allele detected with SRST2 with the same colours as the tree tips. White NA, red P_N_ and black P_O_.

The distribution of *sadP* genes in clinical and non-clinical strains analyzed with core genome phylogeny analysis is shown in Figure 1C. The gene of *sadP* type P_N_ was found in 91.9 % of systemic *S. suis* isolates (88.1 % of strains isolated from brain), whereas only less than 1 % of strains had the gene encoding *sadP* type P_O._ Based on the maximum-likelihood core genome phylogeny analysis the type P_N_ SadP gene is found in clonal virulent *S. suis* strains, which mostly belong to serotype 2. Interestingly, also serotype 14 strains isolated from human meningitis cases have type P_N_ gene. Type P_O_ *sadP* was found more frequently in respiratory and non-clinical strains.

### Purification and analysis of glycosphingolipid composition of porcine lung

The interaction of recombinant SadP with globo series glycolipids has not been thoroughly characterized. Moreover, whether SadP binds to Gb3 or Gb4 isolated from the pig tissues has not yet been directly shown. The glycolipids were isolated from porcine lung and erythrocytes, in order to control that the glycolipids isolated from the lungs were not contaminants derived from the erythrocytes remaining in the lungs. The total non-acid glycosphingolipid fractions were hydrolyzed with endoglycoceramidase II of *Rhodococcus* sp., and the oligosaccharides thereby obtained were analyzed by LC-ESI/MS using graphitized carbon columns. This method gives resolution of isomeric oligosaccharides, and the MS^2^ analyses gives complete sequence information and allows differentiation of linkage positions by diagnostic cross-ring ^0,2^A-type fragment ions (20).

The base peak chromatograms from LC-ESI/MS of the oligosaccharides obtained by hydrolysis of the total non-acid glycosphingolipid fractions of porcine erythrocytes and lung are shown in Figure 2A. By comparison of the retention times and MS^2^ spectra of oligosaccharides obtained from reference glycosphingolipids, the major oligosaccharides obtained from porcine lung were tentatively identified as globotriaosylceramide (Gb3, detected as a [M-H^+^]^−^ ion at *m/z* 503) and the blood group H type 2 pentasaccharide (H5-2, detected as a [M-H^+^]^−^ ion at *m/z* 852) (Figure 2A). The major oligosaccharides from porcine erythrocytes were globotriaosylceramide (Gb3, detected as a [M-H^+^]^−^ ion at *m/z* 503), globotetraosylceramide (Gb4, detected as a [M-H^+^]^−^ ion at *m/z* 706), and the blood group A type 2 hexasaccharide (A6-2, detected as a [M-H^+^]^−^ ion at *m/z* 1055) (Fig 2B). Thus, globotriaosylceramide was found in both samples, but otherwise the two samples were significantly different. The total non-acid glycosphingolipid fraction from porcine lung was thereafter separated by Iatrobeads chromatography. Aliquots of the fractions obtained were analyzed by thin-layer chromatography, and fractions that were coloured green by anisaldehyde were tested for binding of SadP using the chromatogram binding assay. The fractions were thereafter pooled according to the mobility on thin-layer chromatograms and their SadP binding activity.

**Fig 2.**
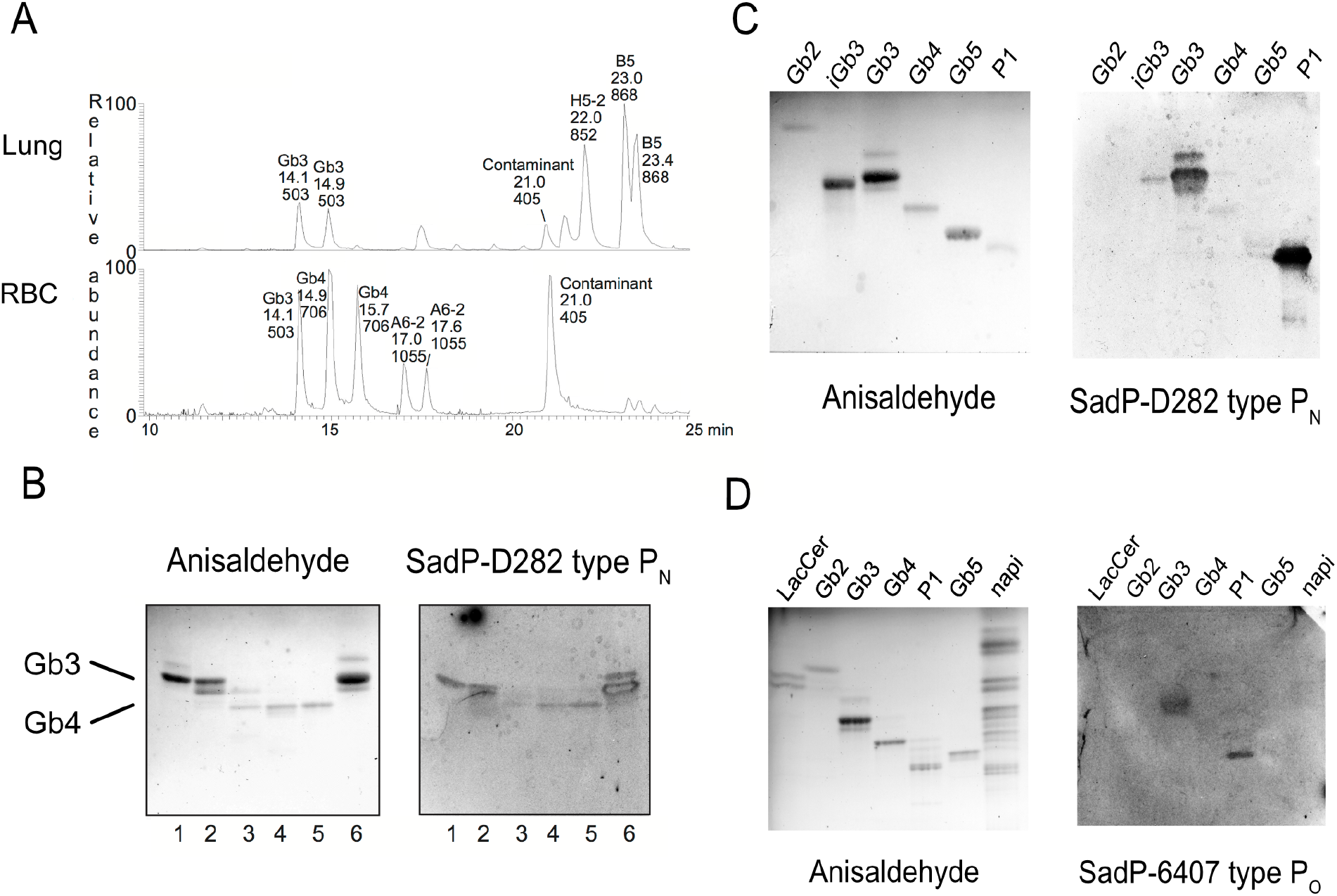
Different glycolipid binding specificities of SadP type P_N_ from systemic *S. suis* strains and type P_O_ SadP from respiratory strains towards pig lung glycolipids and purified globo series glycolipids. **A. Comparison of the total non-acid glycosphingolipid fractions from porcine lung and erythrocytes.** Base peak chromatogram from LC-ESI/MS of the oligosaccharides derived the total non-acid glycosphingolipid fraction from porcine lung by digestion with *Rhodococcus* endoglycoceramidase II. B. Base peak chromatogram from LC-ESI/MS of the oligosaccharides derived the total non-acid glycosphingolipid fraction from porcine erythrocytes by digestion with *Rhodococcus* endoglycoceramidase II. The identification of individual glycosphingolipid-derived oligosaccharides was based on their determined molecular masses and subsequent MS^2^ sequencing. The oligosaccharides identified in the chromatograms are: Gb3, Galα1–4Galβ1– 4Glc; H5-2, Fucα2–3Galβ1–4GlcNAcβ1–3Galβ1–4Glc; B5, Galα1–3Galβ1–4GlcNAcβ1–3Galβ1–4Glc; Gb4, GalNAcβ1–3Galα1–4Galβ1–4Glc; A6-2, GalNAcα1–3(Fucα2–3)Galβ1–4GlcNAcβ1–3Galβ1–4Glc. **B. SadP binding tri- and tetraglycosylceramides isolated from porcine lung.** Thin-layer chromatogram after detection with anisaldehyde (A), and autoradiograms obtained by binding of SadP (B), followed by autoradiography for 12 h, as described under “Methods”. The solvent system used was chloroform/methanol/water (60:35:8, by volume). The lanes were: Lane 1, fraction PL-1 isolated from pig lung, 1 μg; Lane 2, fraction PL-2 isolated from pig lung, 1 μg; Lane 3, fraction PL-3 isolated from pig lung, 1 μg; Lane 4, fraction PL-4 isolated from pig lung, 1 μg; Lane 5, reference globotetraosylceramide (GalNAcβ1– 3Galα1–4Galβ1–4Glcβ1Cer), 2 μg; Lane 6, reference globotriaosylceramide (Galα1–4Galβ1–4Glcβ1Cer), 2 μg. **C. TLC overlay assay with purified glycolipids**, **SadP-D282 type P_N_** 1. Gb2, galabiaosylceramide 2 μg, 2. iGb3, isoglotriosylceramide 4 μg, 3 Gb3, globotriosylceramide 4 μg, 4. Gb4, globotetraosylceramide 4 μg, 5. Gb5, Forssman 4 μg, 6. P1 pentaosylceramide 4 μg. **D. SadP-6407 type P_O_** 1. Lactosylceramide 4 μg, 2. Galabiaosylceramide 4 μg, 3. Globotriaosylceramide 4 μg, 4. Globotetraosylceramide 4 μg, 5. P1, pentaosylceramide 4 μg, 6. Forssman pentaosylceramide 4 μg, 7. Non-acid glycosphingolipids of porcine small intestine (blood group O) 40 μg.

### The binding of SadP to neutral glycolipids isolated from porcine lung using TLC overlay assay

The separations gave two SadP binding fractions migrating in the triglycosylceramide region. One fraction (0.2 mg) migrating as a single band was denoted fraction PL-1 (Fig 2B, lane 1), and one fraction (0.3 mg) migrating as a double band was denoted fraction PL-2 (Fig 2B, lane 2). In addition, one SadP binding fraction (0.4 mg) with compounds migrating as tetraglycosylceramides was obtained (denoted fraction PL-4; Fig 2B, lane 4).

### LC-ESI /MS of native SadP binding glycosphingolipids from porcine lung

ESI/MS of the native fraction PL-1 gave a major [M-H^+^]^−^ ion at *m/z* 1132 (S1A Fig), indicating a glycosphingolipid with three Hex, and sphingosine with non-hydroxy 24:1 fatty acid. MS^2^ of the [M-H^+^]^−^ ion at *m/z* 1132 gave a series of Y ions (Y_0_ at *m/z* 646, Y_1_ at *m/z* 808, and Y_2_ at *m/z* 970) demonstrating a Hex-Hex-Hex sequence combined with sphingosine with non-hydroxy 24:1 fatty acid (S1B Fig).

A [M-H^+^]^−^ ion at *m/z* 1132 was also obtained by ESI/MS of the native fraction PL-2 (S2 Fig). Here, the major [M-H^+^]^−^ ion was seen at *m/z* 1022 (S2AB Fig), demonstrating a glycosphingolipid with three Hex, and sphingosine with non-hydroxy 16:0 fatty acid. Also, here a Hex-Hex-Hex sequence was demonstrated by the series of Y ions (Y_0_ at *m/z* 536, Y_1_ at *m/z* 698, and Y_2_ at *m/z* 860) from MS^2^ (S2D Fig).

### Endoglycoceramidase digestion and LC-ESI /MS of SadP binding glycosphingolipids from porcine lung

The base peak chromatograms from LC-ESI/MS of the oligosaccharides obtained by endoglycoceramidase digestion of fractions PL-1 and PL-2 were very similar (S3AB Fig). Both had a [M-H^+^]^−^ ion at *m/z* 503 which eluted at the same retention time (17.4-18.4 min) as the saccharide obtained from reference globotriaosylceramide (S3C Fig), while the saccharide from reference isoglobotriaosylceramide eluted at 20.0-20.4 min (Figure S3D).

MS^2^ of the ion at *m/z* 503 of fractions PL-1 and PL-2 gave in both cases two C-type fragment ions (C_1_ at *m/z* 179 and C_2_ at *m/z* 341) identifying a Hex-Hex-Hex sequence (Figure S3EF). A 4-substitution of the internal Hex was demonstrated by the ^0,2^A_2_ fragment ion at *m/z* 281 (20–22). Taken together with the similarity to the MS^2^ spectrum of reference globotriaosyl saccharide (S3G Fig), this allowed identification of the saccharides of fractions PL-1 and PL-2 as globotriaosyl saccharides (Galα1–4Galβ4Glc).

LC-ESI/MS of the oligosaccharides obtained by hydrolysis of fraction PL-4 with *Rhodococcus* endoglycoceramidase II allowed a tentative identification of a globotetraosyl saccharide (GalNAcβ1–3Galα1– 4Galβ1–4Glc). This conclusion was based on the following spectral features: First, the base peak chromatogram of fraction PL-4 had a [M-H^+^]^−^ ion at *m/z* 706 (Figure S4A), and MS^2^ of this ion (S4B Fig) gave a C-type fragment ion series (C_1_ at *m/z* 220, C_2_ at *m/z* 382, and C_3_ at *m/z* 544), demonstrating a HexNAc-Hex-Hex-Hex sequence. The ^0,2^A_3_ fragment ion at *m/z* 484 demonstrated a 4-substituted Hex, while the ^0,2^A_4_ ion at *m/z* 646, and the ^0,2^A_4_-H_2_O ion at *m/z* 628, were derived from cross-ring cleavage of the 4-substituted Glc of the lactose unit at the reducing end. The features of this MS^2^ spectrum were very similar to the MS^2^ spectrum of the reference globotetraosyl saccharide (S4C Fig)

The base peak chromatogram of fraction PL-4 (S4A Fig) also had two [M-H^+^]^−^ ions at *m/z* 998, eluting at 19.4 min and 21.4 min. In both cases MS^2^ demonstrated a Fuc.Hex-(Fuc-)HexNAc-Hex-Hex sequence, and the diagnostic ion at *m/z* 348 (S4D Fig) identified a Le^b^ hexasaccharide, whereas the diagnostic ion at *m/z* 510 (S4E Fig) identified a Le^y^ hexasaccharide (21).

### SadP subtype P_N_ and P_O_ binding to purified glycolipids

TLC overlay assay is a powerful method to analyze glycolipid receptor function. The carbohydrate-binding specificity of SadP to globo series isoreceptors was determined using purified glycolipids (all TLC binding results are summarized in Table 1). Type P_N_ bound preferentially to Gb3 and P1 glycolipid, and also weakly to Gb4 and the Forssman glycolipid (Fig 2C). SadP type P_O_ also bound strongly to Gb3 and P1, i.e. to glycolipids with terminal Galα1–4Gal, whereas there was no binding to Gb4 and the Forssman glycolipid (Fig 2D). There was no binding to the non-acid glycolipid fraction from the pig intestine, containing the Globo H.

**Table 1.**
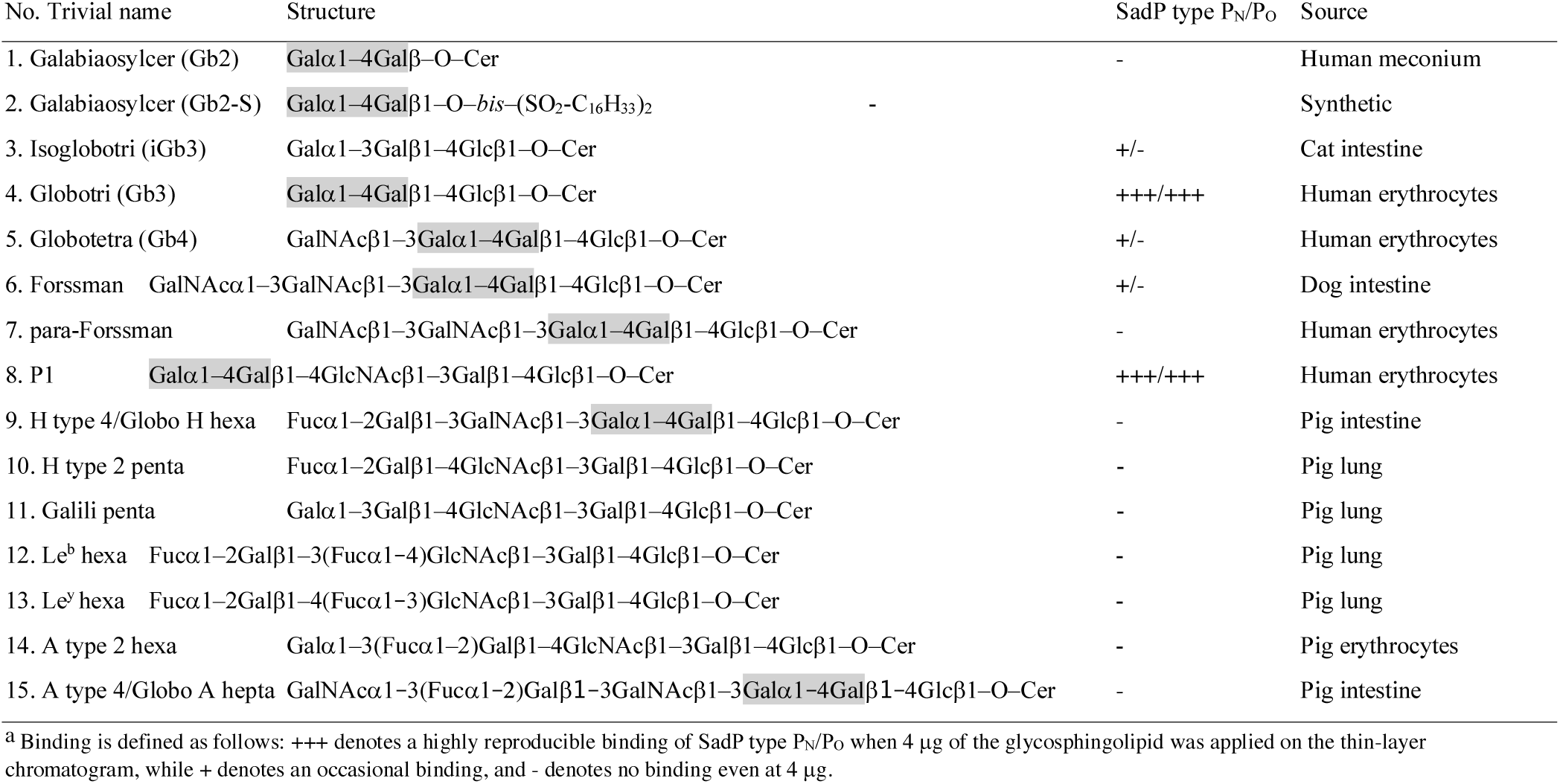
Binding of SadP to glycosphingolipids on thin-layer chromatograms.

The type P_N_ SadP binding to dilutions of globo series glycolipids was analyzed with TLC overlay assay to the detection limit for the various glycolipids (S5A Fig). The full length adhesin bound mainly to P1 glycolipid, with a detection limit at 0.4 μg, whereas the detection limit for the Gb3 glycolipid was 0.8 μg. The detection limit of SadP(31-328) *N*-terminal domain was 0.2 μg for both Gb3 and P1.

Next, the SadP binding to 250 ng of glycolipids immobilized in plastic microtiter wells was quantitatively evaluated by solid phase binding assay (S5B Fig). No binding to the Gb2 glycolipid, galabiosylceramide was detected, whereas there was weak binding to Galα1–4Galβ1–O-*bis-*(SO_2_-C_16_H_33_)_2_. SadP was found to bind stronger to Gb3 with non-hydroxy ceramide than to Gb3 with hydroxy ceramide. There was also binding to Gb4 and iGb3 isoglobotriaosylceramide. Binding to P1 pentaosylceramide was at the same level as binding to Gb3 with non-hydroxy ceramide. The difference in the binding to iGb3 (Galα1– 3Gal-containing glycolipid) in the microtiter well assay compared to TLC overlay assay could be due to the difference in the presentation of the binding epitope and the clustering of the glycolipid moieties.

### Galα1–4Gal-dependent endothelial cell binding activity of *S. suis* SadP

*S. suis* type P_N_ WT strain D282 and the corresponding mutant strain D282ΔsadP, containing an insertion mutation into SadP gene, were compared for their binding activity to cultured cell line EA.hy926, which is a primary human umbilical vein cell line fused with a thioguanine-resistant clone of A549 (human lung adenocarcinoma cell line). In addition, binding of purified fluorescently labelled SadP was analysed for the binding to EA.hy926 cells. The cells were grown into glass coverslips for 48 h and the bacteria grown into early log phase were added to cells and were let to adhere for 1 h. After washing and staining with DiffQuick the bacteria were enumerated. The average adherence of wild-type bacteria was calculated to be 8.7 ± 0.5 bacteria/field, whereas the mean adhesion of 1.3 ± 0.5 of the insertion mutant ΔsadP was significantly less (unpaired t test > 0.0001) (Fig 3A). In addition, the binding of the WT strain was inhibited with 10 μg/ml pigeon ovomucoid, which contains *N*-linked glycans with terminal Galα1–4Gal (23). The inhibitor reduced the adherence of *S. suis* bacteria to the cells to 0.7 ± 0.1 bacteria/field.

**Fig 3.**
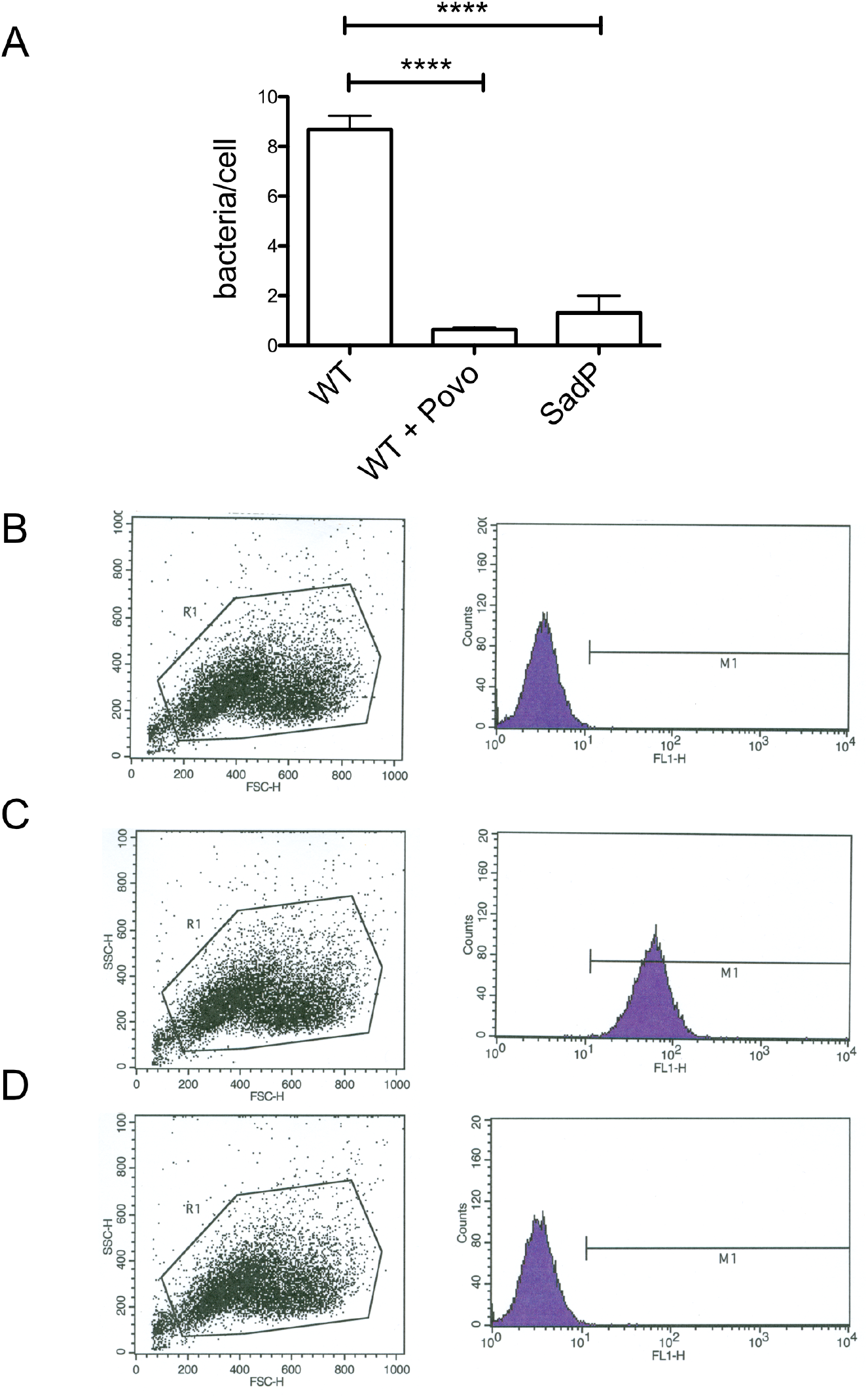
SadP binding to EA.hy926 cell line. **A.** Binding of S. suis D282 wild type serotype 2 strain and the SadP insertion mutation to EA.hy926 cells grown on coverslips. The binding of the wild type bacteria was inhibited with 10 μg/ml of pigeon ovomucoid. Flow cytometry analysis of SadP(31-328) type P_N_ binding to EA.hy926 cells. **B.** Buffer control. **C.** SadP adhesin (0.4 μg/ml) and **D.** SadP adhesin in the presence of 10 μg/ml of pigeon ovomucoid.

For flow cytometry assay, the cells were grown in wells, washed and tested for the binding of FITC labelled SadP. After washing, the cells were harvested with scraping and analyzed with flow cytometer. The flow cytometry results (dot blot and the histogram) are presented in Figure 3BCD. The median SadP binding with and without inhibitor (pigeon ovomucoid, 10 μg/ml) were 56.2 and 14.9 per 10 000 total events (median of untreated cells was 12.8).

### Fine specificities toward Gb3 and Gb4 oligosaccharide structures

Next, the ability of the type P_N_ SadP-D282(31-328) and P_O_ SadP-6407(31-328) recombinant adhesin constructs to recognize galabiose glycoconjugates was analyzed using isothermal titration calorimetry (Fig 4AB). Both type P_N_ and P_O_ adhesins were titrated with oligosaccharides representing the terminal epitopes of Gb3 and Gb4, the TMSEt (2-trimethylsilylethyl) glycosides of Galα1–4Gal and GalNAcβ1–3Galα1–4Gal (24). The dissociation constant values (K_D_) of type P_N_ SadP-D282(31-328) and type P_O_ SadP-6407(31-328) interaction with Galα1–4Gal were 13.0 ± 1.5 μM (n=0.85, ΔH=−69.0±2.0 and -TΔS=41.1) and 3.5 ± 1.6 μM (n=0.85, ΔH=−40.0±2.7 and -TΔS=8.8) respectively. The K_D_ for interaction of P_N_ and P_O_ adhesins with GalNAcβ1–3Galα1–4Gal compared to Galα1–4Gal were 2.6-fold higher with type P_N_ (K_D_= 33.7±3.5 μM, n=1.14, ΔH=−40.1±2.0 and -TΔS=14.0) and 269-higher with type P_O_ (K_D_= 940±680 μM, n=0.13, ΔH=−197±5500 and -TΔS=179).

**Fig 4.**
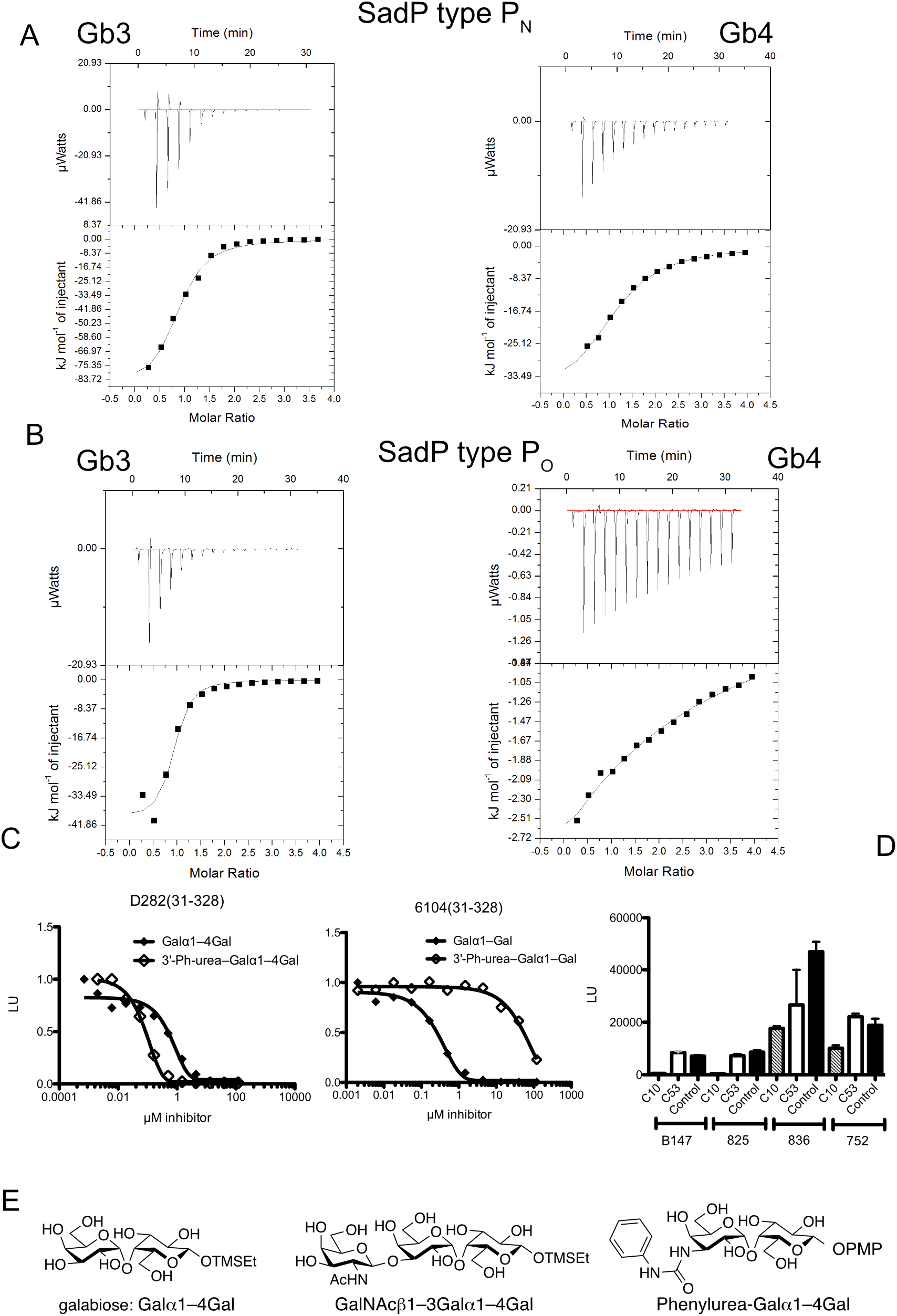
Oligosaccharide specificity of SadP from systemic (type P_N_) and respiratory (type P_O_) *S. suis* strains analyzed with glycans representing the terminal saccharide structure of Gb3 and Gb4. **A. and B.** ITC of SadP type P_N_ and type P_O_ with Galα1–4Galβ1– and GalNAcβ1–3Galα1–4Galβ1– TMSEt derivatives **C.** AlphaScreen inhibition assay with Galα1–4Galβ1– and 3’Phenylurea-Galα1–4Galβ1– methoxyphenyl (-PMP = 4-methoxyphenyl). **D**. AlphaScreen inhibition assay of Galα1–4Galβ1– (compound C10) and 3’Phenylurea-Galα1–4Galβ1– methoxyphenyl (compound C53) with recombinant N-terminal domains cloned from 4 different type P_O_ *S. suis* strains **E**. Structures of the ligands in this study.

Then, both adhesins were analyzed with AlphaScreen competitive inhibition assay (Fig 4C) for their interaction with Galα1–4Gal in comparison to a glycomimetic inhibitor 3’-phenylurea-derivative (3’-phenylurea-Galα1–4Gal β1–methoxyphenyl), which is a low nanomolar inhibitor of type P_N_ SadP (19, 25). The binding of His-tagged recombinant adhesins to biotinylated pigeon ovomucoid was inhibited with the above oligosaccharides. The IC_50_ values of type P_N_ adhesin for Galα1–4Galβ1–methoxyphenyl and 3’-phenylurea-Galα1–4Gal β1–methoxyphenyl were 1 μM and 0.030 μM, whereas with type P_O_ adhesin the corresponding values were 0.3 μM and 70 μM respectively. Also 4 other recombinant type P_O_ SadP N-terminal-domains were tested using 10 μM inhibitor concentrations (Fig 4D). For all type P_O_ adhesins the inhibition obtained with the C3’-phenylurea-derivative was weaker than the inhibition obtained with Galα1– 4Gal.

### Site-specific mutation changes the specificity of type P_N_ recombinant adhesin to type P_O_

The binding mechanism of type P_N_ and P_O_ SadP was further studied by site-directed mutagenesis. Type P_N_ adhesin SadP-D282(125-329) was mutated and the mutants were analyzed for the binding to galabiose oligosaccharides with AlphaScreen competitive inhibition assay and ITC. Multiple alignment of SadP homologues allowed us to predict amino acids from the galabiose-binding region that differed between type P_N_ and P_O_ (Fig 1A). Based on that, the site-specific amino acid change N285D and E292Q, deletion Δ244-246 and N285D&Δ244-246 mutants were constructed into the type P_N_ SadP(125-328) cloned into the plasmid pET28a. In addition, the 28 amino acid long region H216-D246 of the type P_N_ was replaced by the corresponding region of type P_O_.

The WT SadP and mutants were expressed and purified as described in the methods and analysed with SDS-PAGE (S6 Fig). The galabiose-binding strengths of the above mutant constructs, full length SadP, SadP *N*-terminal constructs SadP(31-328), SadP(125-328) and as a negative control site-specific mutant SadP(31-328)W258A, were compared by AlphaScreen asssay (S7 Fig). The binding of full-length SadP to biotinylated pigeon ovomucoid was weaker compared to the *N*-terminal domain constructs, which could be due to a larger size of the full length SadP. The size of the protein could increase the distance of the acceptor and donor beads, and hence reduce the transmission of the singlet oxygen during the assay (19). As a negative control for the assay, there was no binding of site-specific mutant W258A to pigeon ovomucoid. Of the mutants in the type P_N_ SadP(125-328) background, mutants N285D and E292Q bound well whereas the binding of mutants Δ244-246 and H216-D246delinsDELFNRFPNKVDSTNNGDGAPFRFFNKE was reduced. The binding of N285D&Δ244-246 mutant was abolished.

Next the specificities of mutants were analyzed with AlphaScreen assay using phenylurea-derivative and Galα1–4Gal (S7B - F Figs). All mutants were equally inhibited by the Galα1–4Gal oligosaccharide, but only with the site-specific mutant N285D the inhibitory power of Phenylurea-Galα1–4Gal was weaker compared to Galα1–4Gal, suggesting a specific switch from type P_N_ to P_O_ specificity.

Next, the Alphascreen inhibition assay was used to compare the effect of site-specific mutation in N285D to the specificity between Galα1–4Gal (terminal Gb3 structure) and GalNAcβ1–3Galα1–4Gal (terminal Gb4 structure) TMSEt glycosides. SadP(125-329) type P_N_ and site-specific mutant N285D were similarly inhibited with Galα1–4Gal, whereas N285D was not inhibited with even higher concentrations of GalNAcβ1–3Galα1–4Gal (Fig 5AB).

**Fig 5.**
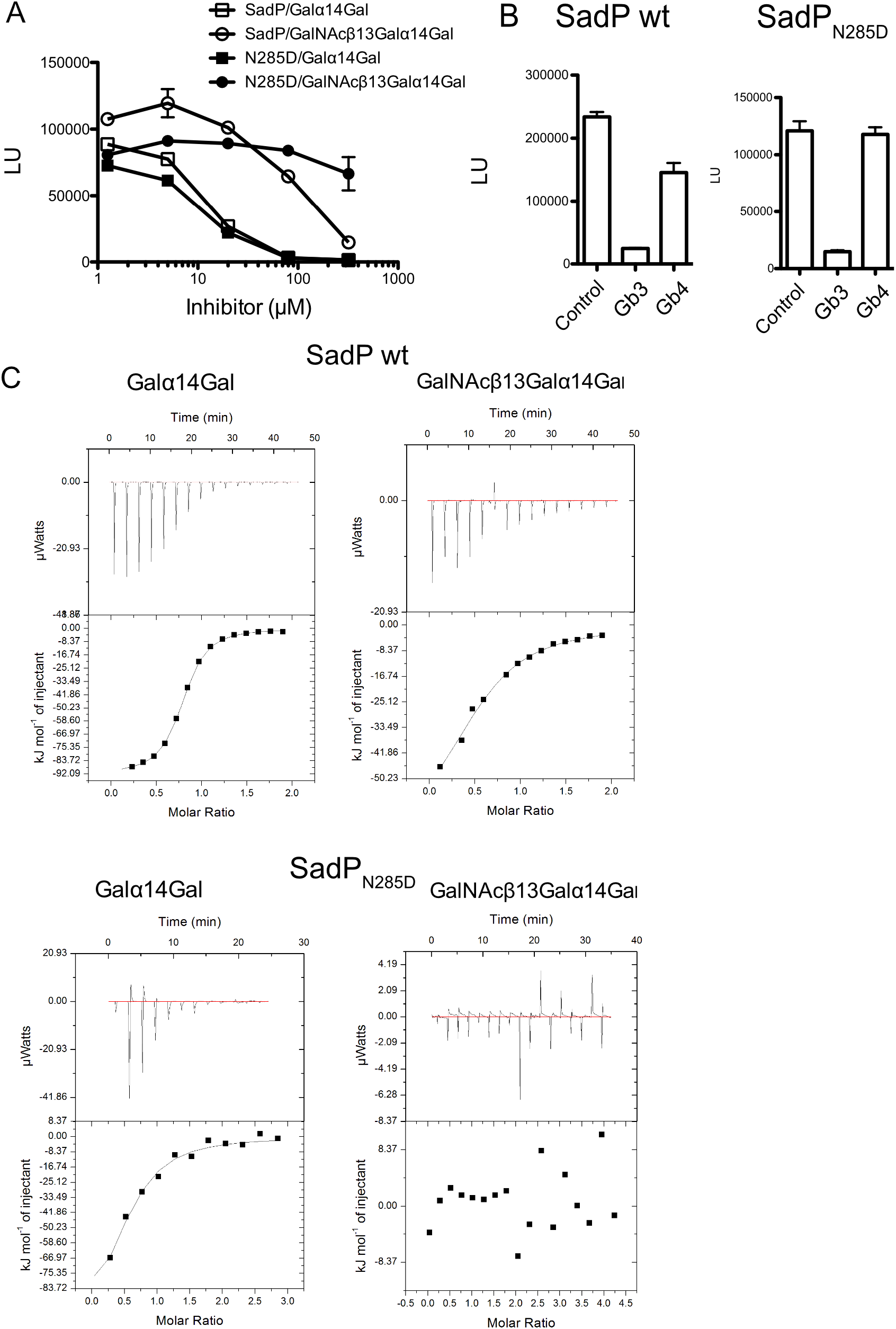
A. Single amino acid site-specific mutation N285D of SadP from systemic strains changes its Gb4 binding specificity to type P_O_ Gb3 binding specificity of respiratory strains. **A.** Inhibition of WT SadP(125-329) and its site-specific mutant SadP(125-329)N285D with the dilutions series (1.25 to 320 μM, duplicate determinations) of TMSEt (2-trimethylsilylethyl) glycosides of Galα1–4Gal and GalNAcβ1– 3Galα1–4Gal representing the Gb3 and Gb4 oligosaccharides. **B.** Inhibition assay with 5 μM of Galα1–4Gal and GalNAcβ1–3Galα1–4Gal TMSEt glycosides. **C.** ITC measurements of WT SadP and N285D mutant with the TMSEt glycosides of Galα1–4Gal and GalNAcβ1–3Galα1–4Gal representing the Gb3 and Gb4 oligosaccharides.

The K_D_ values of type P_N_ SadP(125-328) and site-specific mutant N285D for oligosaccharides was analyzed with isothermal titration calorimetry (Fig 5C). The wild type SadP(125-329) K_D_ values for Galα1– 4Gal and GalNAcβ1–3Galα1–4Gal were 3.4±0.09 (n=0.76, ΔH=92.0±0.3 and -TΔS=60.8) and 36.0±4.8 μM (n=0.56, ΔH=70.2±5.0 and -TΔS=44.7) respectively. The K_D_ of site specific mutant N285D with Galα1–4Gal was 22.7±7.2 μM (n=0.52, ΔH=26.8±5.4 and -TΔS=20.5) (6-fold increase compared to wild-type SadP), whereas there was no interaction between N285D and GalNAcβ1–3Galα1–4Gal. Thus, the above results suggest that N285 has a specific role in SadP type P_N_ binding to GalNAcβ1–3Galα1–4Gal.

The effect of N285D mutation to SadP binding to glycolipids incorporated into POPC and POPC-cholesterol liposomes was analyzed by dotting liposomes to PVDF membrane and binding of his-tagged adhesins to immobilized liposomes (Fig 6). WT SadP type P_N_ bound to Gb4 of both non-cholesterol and cholesterol-containing liposomes, whereas there was no binding by N285D mutant. The acyl chain length or hydroxylation of ceramide had no major effects for Gb3 binding.

**Fig 6.**
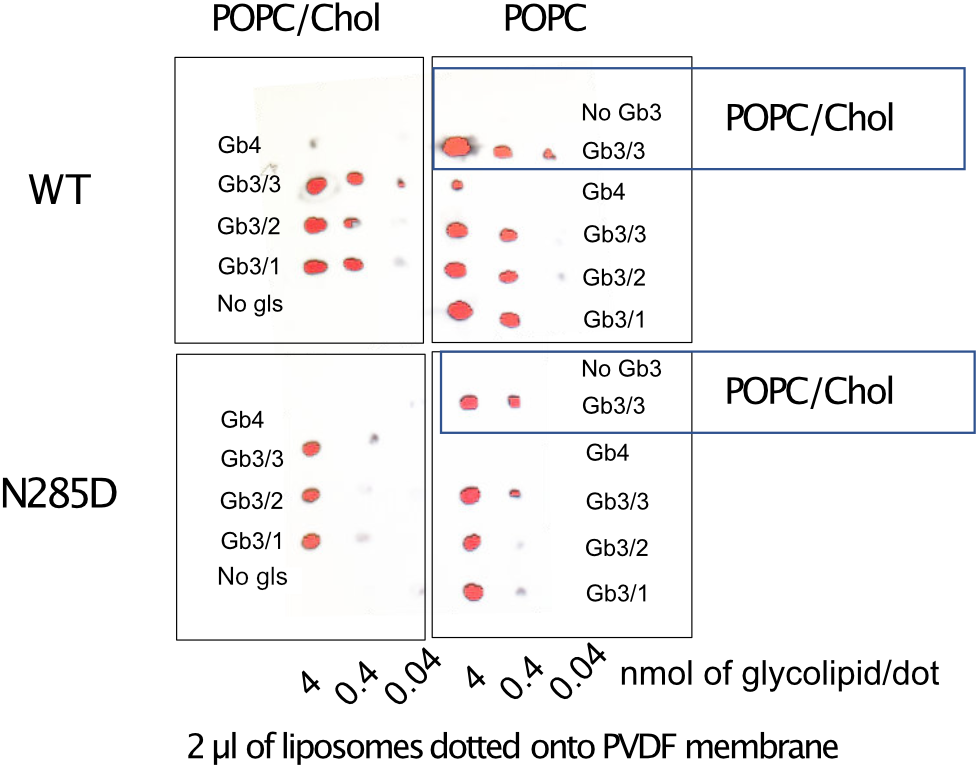
SadP and SadP_N285D_ binding to liposomes. POPC/glycolipid liposomes (4, 0.4 and 0.04 nmol/dot) and POPC/cholesterol/globosylceramide liposomes (4, 0.4 and 0.04 nmol/dot) were applied onto PVDF membrane. WT, His-tagged SadP(125-329) type P_N_; N285D, site-specific mutant of SadP(125-329); Replicate membranes containing dotted liposomes with or without cholesterol (see Methods) were probed with **SadP and SadP_N285D_**. The bound proteins were detected with anti-His and HRP-labelled antibodies as described in the methods and membrane was imaged with FujiLAS-4000. The globo series glycolipids used were: Gb4, d18:1-24:0 + d18:1 – 16:0; Gb3/1, Gb3 d18:1-24:0; Gb3/2, Gb3 d18:1-16:0; Gb3/3, Gb3 d18:1-h16:0.

### Structural comparison of recombinant SadP type P_N_ and its N285D mutant

The site-specific mutant SadP(125-329)N285D was crystallized with five molecules in the crystallographic asymmetric unit (Fig 7A, Table S1). All molecules are similar with subtle differences as suggested by the low root mean square deviation (r.m.s.d) between them (0.25-0.31 Å). The structure of the N285D mutant is similar to that of the native SadP (PDB id 5BOB) with r.m.s.d. of 0.26 Å for 199 aligned residues. The structure consists of three α-helixes and ten β-strands (β1-β10) that form a β-sandwich core domain (Fig 7B). The first β-sheet of the β-sandwich is formed by antiparallel β1-β10-β9-β4-β7-β6 strands and the second β-sheet by β2-β3-β8-β5 in antiparallel fashion.

**Fig 7.**
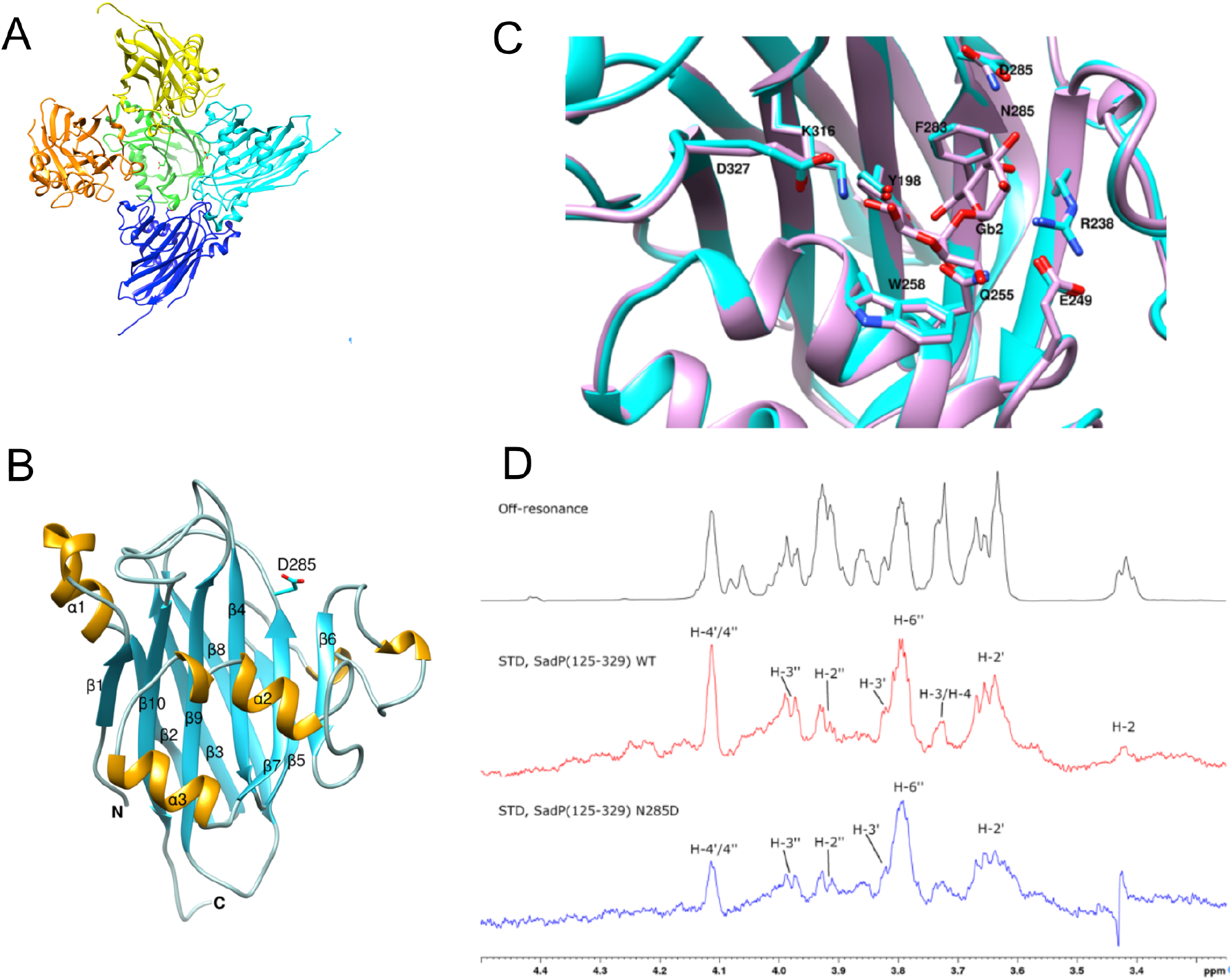
**A.** Arrangement of the five site-specifically mutated SadP(125-329)N285D molecules in the crystallographic asymmetric unit. Each molecule is shown as ribbon and colored differently. **B.** Ribbon diagram of SadP(125-329)N285D structure. α-Helices are shown in yellow color and β-strands in cyan. D285 is depicted as sticks. **C.** Structural superposition of Galα1–4Gal-bound Fhc (PDB 5BOA) structure (identical to SadP type P_N_ (pink) onto SadP(125-329)_N285D_ (cyan). Residues involved in binding interactions are shown in stick representation. **D.** STD-NMR of binding of wild type SadP(125-329) type P_N_ and its site-specific mutant N285D. Peaks in the STD spectra indicate hydrogens (C-H) that are in close proximity to the protein.

Structural superposition with the Galα1–4Gal – Fhc structure revealed small differences (Fig 7C). The side chain of D285 was found slightly rotated compared to the N285 side chain in the wild type.

### Resolution of the interaction mechanisms of SadP to globo series glycolipids and C3’-Phenylurea–Galα1–4Gal

The key amino acids of both types P_N_ and P_O_ are conserved. Y198, Q255, R238 and G233 forms hydrogen bonds with the HO–3’, HO-4’ and HO-6’ of the α’-Gal and the K316 with HO-2 and HO-3 of the β-Gal pyranose ring. W258 most likely forms CH-π interactions with the hydrophobic face of the disaccharide (Fig 7C). Type P_N_ SadP N285 is on the edge of the binding pocket, and it potentially interacts with the terminal GalNAcβ1–3 of Gb4 or 3-phenylurea-group. To study the SadP interaction with the receptor saccharide Galα1–4Galβ1–4Glc in solution, STD-NMR was applied using type P_N_ SadP(125-329) and the corresponding mutant N285D (Fig 7D). STD-NMR results suggest that both the type P_N_ SadP and the mutant N285D interact mainly with the two terminal galactose units of Gb3. With the mutant N285D the interactions of H-4ʹ and H-4ʹʹ are, however, somewhat weaker compared to the interaction of H-6ʹʹ. Additionally, the mutant N285D does not seem to have a clear interaction with the β-glucoside unit at the reducing end of Gb3, whereas the type P_N_ SadP appears to have interactions with H-2, H-3 and H-4.

In order to find a plausible hypothesis explaining the binding preferences of the galabiose, GalNAcβ1– 3Galα1–4Gal and phenylurea-galabioside for WT and the N285D mutant, molecular dynamic simulations of their respective complexes were performed. Briefly, the ligand structures were built without the TMSEt aglycon to simplify the simulations and were then placed in the binding site of SadP (pdb id 5BOA) with the Galα1–4Gal disaccharide oriented as in the crystal structure. The terminal GalNAcβ1 residue was oriented with the dihedral glycosidic bond angles minimized with the H1 parallel to H4’ of Galα1–4Gal. The three complexes with WT were mutated by exchanging N285 for D285. The complexes were unrestrainedly subjected to 100 ns molecular dynamics simulations and all starting conformations converged towards similar stable complex geometries (Fig 8A-E), except for the phenylurea derivative in complex with the N285D mutant for which light constraints on the backbone atoms of protein secondary structures were applied (Fig 8F). The α-face of the β-galactoside residue of all three ligands stacked to W258 forming CH-π interactions and the galabiose disaccharide moiety in all simulations retained the interactions and hydrogen bonds observed in the X-ray structure with the Gb3 trisaccharide (PDB id 5boa). The HO-4’ of galabiose did not interact directly with N285, but instead via N285 sidechain NH hydrogen bond donation to a network of hydrogen bonds involving 3-5 water molecules and the G233 carbonyl oxygen (Fig 8A and S9 movie). Mutation of N285 to D285 influenced the directionalities of the hydrogen bonding partners in this network (Fig 8D and S10 movie), which may explain the different STD-NMR data with WT and the D285 mutant. The simulations with the GalNAcβ1–3Galα1–4Gal and the phenylurea-galabioside in complex with the WT protein converged to geometries that revealed a highly populated interaction between the ligands and N285. The ring oxygen of the GalNAc residue of the trisaccharide and the urea carbonyl oxygen of the phenylurea derivative accepted hydrogen bonds from the N285 side chain NH_2_ (Fig 8B-C and S11-S12 movies). The GalNAc HO-4 of the trisaccharide formed transient hydrogen bonds to side chain amide groups of E286 and E234, while the phenyl ring of the phenylurea ligand did not form any persistent interactions with any protein residue. The highly populated hydrogen bonds of the trisaccharide GalNAc ring oxygen and the phenylurea carbonyl oxygen to N285 could not form in the N285D mutant (Fig E-F and S13-14 movies). Instead, it was disrupted by the formation of a water network between the D285 side chain carboxylate and the GalNAc ring oxygen of the trisaccharide (Fig 8E), while the phenylurea moiety of the phenylurea galabioside rotated away from the D285 side chain carboxylate (Fig 8F). This lack of direct hydrogen bond interactions of the GalNAc ring oxygen and urea carbonyl oxygen may at least partly explain the less efficient binding of the trisaccharide and phenyl urea derivative to the N285D mutant.

**Fig 8.**
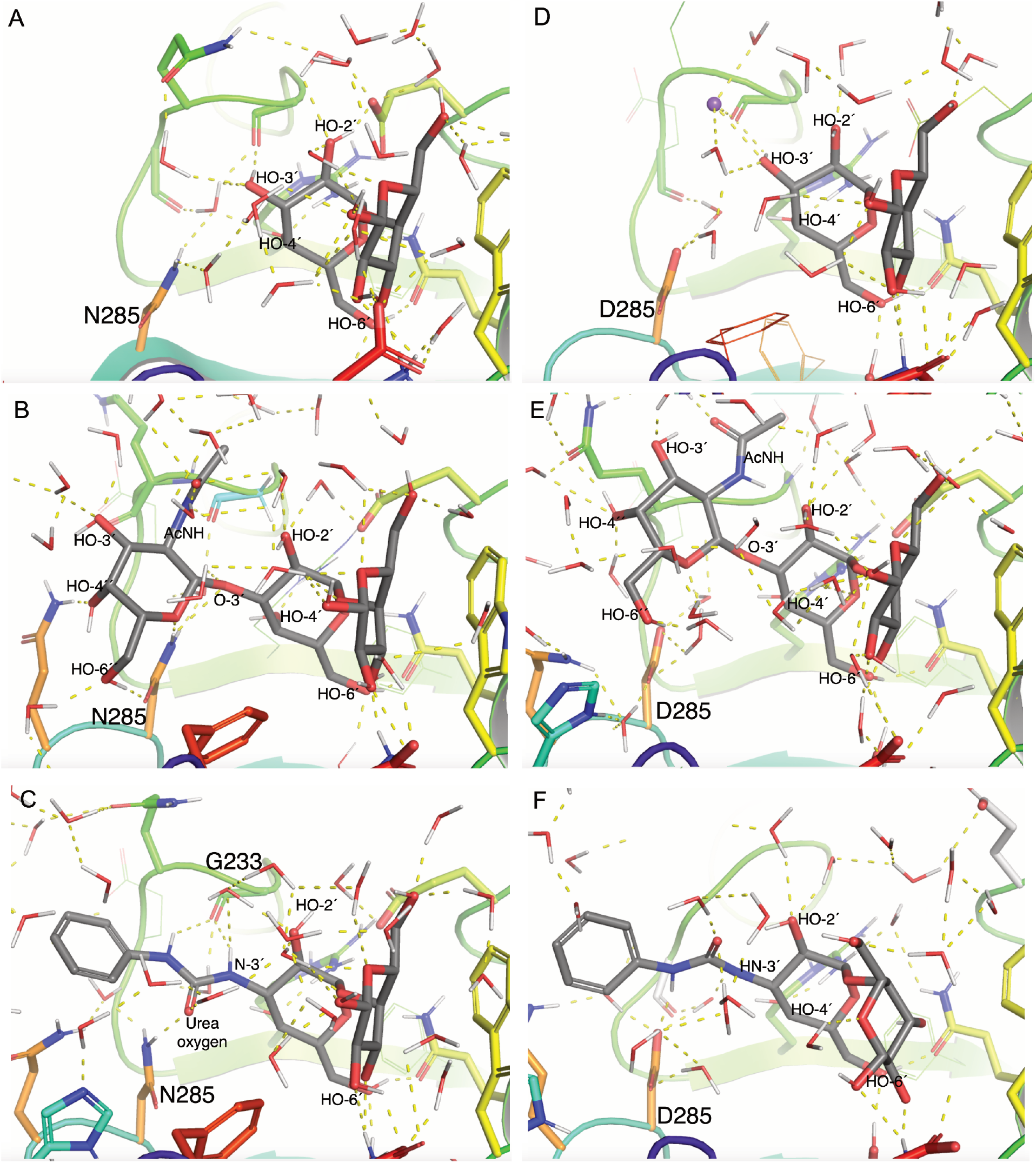
Molecular dynamics of SadP type P_N_ binding to Gb4 saccharide and glycomimetic phenylurea-galabiose inhibitor. Representative MD snapshots (at 90 ns) of SadP. WT in complex with **A**. Galα1– 4Gal, **B**. GalNAcβ1–3Galα1–4Galβ, and **C.** C3′-phenylurea-Galα1–4Galβ. Representative MD snapshots of SadP N285D in complex with **D.** Galα1–4Gal, **E.** GalNAcβ1–3Galα1–4Galβ, and **F.** C3′-phenylurea-Galα1– 4Galβ.

## Discussion

SadP carbohydrate specificity to neutral glycolipids from lungs was studied by TLC overlay assay. The carbohydrate receptors for *S. suis* adhesion in respiratory tract and lungs have remained elusive. We studied here how SadP type P_N_ and P_O_ recognize glycolipids isolated from pig lung. The major pig lung neutral glycolipids were identified as Gb3, Gb4, the H type 2 pentaosylceramide, the Galili pentaosylceramide and Le^b^ and Le^y^ hexaosylceramides (Fig 2). In TLC overlay assay, type P_N_ SadP bound to Gb3 and Gb4, whereas type P_O_ SadP bound only to Gb3. Both SadP types showed no binding to Galα1–3Gal-containing Galili pentaosylceramide. (Fig 2AB, Table 1). Therefore, our findings suggest that globo series glycolipids are the major carbohydrate receptors for *S. suis* in pig lung. Although fucose-binding lectins have been described both in bacterial and fungal lung pathogens, no fucose-specific binding activity was found in SadP. The globo series glycolipids have been shown to be widely expressed in many tissues of pigs (26); hence, they can potentially serve as SadP receptors for other host organs and brain capillary veins critical for *S. suis* invasive infection. We show that the binding of type P_N_ *S. suis* serotype 2 *sadP-*mutant to endothelial cell line was significantly reduced and binding of both the WT *S. suis* bacteria and recombinant SadP adhesin were inhibited with the galabiose-containing pigeon ovomucoid.

The oligosaccharide inhibition assays performed with both whole bacteria and recombinant adhesin suggest that the reducing end β-Glc of Gb3 is required for optimal binding (16, 19). The TLC overlay assay using purified glycolipids showed that SadP bound strongest to the P1 glycolipid (Fig 2BC and S5A Fig). The preferential binding to the P1 glycolipid might also be due to a more optimal presentation of the epitope on the longer core chain, whereas hindered presentation of binding epitope in galabiosyl ceramide might explain, why SadP did not bind to Gb2 in TLC overlay assay. The presence of P1 structure is not well known in pig, however it was found in α1,3GalT knockout pig kidney, since this glycosyltransferase is a direct competitor of α1,4GalT (27). This might be important for surveillance of pathogens in xenotransplantation.

The fine specificity of SadP subtypes was further analyzed with ITC and competitive inhibition assay. The ratio of K_D_ values (Galα1–4Gal / GalNAcβ1–3Galα1–4Gal) for type P_N_ SadP was 0.4 whereas for type P_O_ it was 0.004, which is in accordance to the corresponding hemagglutination inhibition MIC values observed with bacterial hemagglutination assays (16). Moreover, the IC_50_ values for Galα1–4Gal and 3’-phenylurea-Galα1–4Gal are in accordance to previous hemagglutination inhibition MIC concentrations for type P_N_ and P_O_ *S. suis* (25).

The mechanism of how the two SadP subtypes differ in their recognition of Gb3, Gb4 and glycomimetic 3-phenylurea-galabiose has not been studied before. The multiple-sequence alignment of SadP showed that specific amino acids G233, E249 and N285 of type P_N_ in galabiose-binding domain are different compared to type P_O_ SadP (Fig 1A). Of these, only substitution from asparagine to aspartate at position 285 is found in all hemagglutinating type P_O_ adhesins. To analyze the mechanisms how SadP type P_N_ can bind to Gb4 terminal trisaccharide (GalNAcβ1–3Galα1–4Gal) and 3’-phenylurea-galabiose, we designed mutations to type P_N_ SadP and analyzed their binding to galabiose. The SadP type P_N_ region from H216 to D246 was swapped to the corresponding region of type P_O_ (to contain substitution from amino acid G to N at position 233). The binding of this mutant to galabiose was reduced compared to the wild type adhesin (S6A Fig), but was still sufficient to further analyze the oligosaccharide specificity. Deletion Δ244-246 and site-specific mutations N285D and E292Q bound also to galabiose at sufficient level (S6A Fig) allowing testing with 3’-phenylurea-derivative to compare for type-specificity. Of these mutants, only N285D lost the ability to be efficiently inhibited by 3’-phenylurea-derivative (S6B-F Fig), suggesting that N285 has an important role in interacting with the phenylurea-group. The amino acid region 216-246 including G233N substitution have possibly no significant role alone in the interaction with HO-3’ substituted phenylurea-galabiose. However, G233 could still synergistically interact with phenylurea. Therefore, the N285D mutant was chosen for further studies for the type P_N_ binding mechanism. The WT and N285D mutant were compared for their binding to the soluble Gb4 terminal trisaccharide GalNAcβ1–3Galα1–4Gal and to globo series glycolipids Gb3 and Gb4 (Figs 5 and 6). The results show consistently that N285 plays a specific role in Gb4 binding as demonstrated by competitive inhibition assay, ITC and liposome binding assay.

Molecular dynamics calculations were performed with the structures representing type P_N_ SadP(125-329) and its N285D site-specific mutant. Both proteins were analyzed with Galα1–4Gal, GalNAcβ1–3Galα1– 4Gal and 3’-phenylurea-Galα1–4Gal. In accordance to the co-crystal structure (PDB id. 5boa), the galabiose HO-4’, HO-6’, HO-2 and HO-3 were hydrogen bonded to Y198, G233, R238, Q255 and K316. Hydrophobic interaction with the α-face of the β-galactoside stacked to W258 forming CH-π interactions. The above interactions were conserved in WT and N285 mutant. WT SadP N285 formed hydrogen bonds to GalNAc O-5 ring oxygen of GalNAcβ1–3Galα1–4Gal and to urea’s carbonyl group (Fig 8B). The amino acid D285 of site-pecific mutant formed hydrogen bonds with solvent water molecules when simulated with GalNAcβ1– 3Galα1–4Gal and 3’-phenylurea-Galα1–4Gal. These results together with the AlphaScreen and ITC data give us a plausible explanation for the molecular mechanism of how type P_N_ SadP interact with Gb4 terminal GalNAc and with phenylurea-galabiose derivative. In addition, WT SadP amino acid G233 interacts with the terminal GalNAc of Gb4 via hydrogen bond to N-Acetyl-group or to urea’s -NH group (Fig 8C).

SadP does not have any sequence homologs in other galabiose-recognizing proteins, however structurally *E. coli* PapG and SadP galabiose-binding sites share some similarities. They both contain a β-sheet composed of antiparallel β-strands, which are adjacent to the galabiose-binding domain (PDB; *E. coli* 1j8r, *S. suis* 5boa). In SadP, the binding site is formed by the alpha-helix and a non-helix loop, whereas in PapG the galabiose binding site is located in the middle of two non-helix loops. Other galabiose binding proteins such as *P. aeruginosa* LecA (4yw6), *E. coli* verotoxin (1bos) and *Lyophyllum decastes* fungal lectin (4ndv) show different combining site architectures. As previously suggested, the galabiose-binding adhesin SadP from gram-positive and *E. coli* P fimbrial adhesins from gram-negative bacteria are an example of convergent evolution towards binding to the galabiose oligosaccharide. The structural similarities thus confirm this hypothesis.

Hemagglutination positive strains expressing type P_N_ SadP from pig and zoonotic meningitis strains belong to serotype 2 and clonal group CC1 (6, 17). *S. suis* hemagglutination type P_O_ strains were mostly found from serotype 4 and were isolated from lungs of strains causing pneumonia (28). The majority of strains that have the gene encoding type P_N_ SadP belongs to a distinct population consisting of systemic isolates, which were identified in a large BRaTP1T consortium functional genomic study (12). *S. suis* strains that cause human meningitis comprise a separate clade and are thought to have evolved when the pig farming was intensified in 1920s (12). Zoonotic *S. suis* strains as a specific population have been thought to exponentially spread to different geographical areas due to the selection of pigs that are optimal in terms of productivity. The BRaTP1T consortium study of *S. suis* genomic signatures for pig and human infections has suggested that the systemic infections are genetically determined and the genome size of highly virulent strains is reduced and enriched with virulence genes. Since the type P_N_ SadP adhesin gene has not been lost in strains causing systemic infections, it is tempting to speculate that the recognition of Gb4 may play a specific role in invasive infections. In a previous study by Ferrando et al. (2017), type P_N_ SadP (designated SadP1) present in CC1 clonal group of *S. suis* strains (17) was found to specifically adhere better to human intestinal cells (Caco-2) than to pig intestinal cells. This could indicate that type P_N_ SadP has specific binding preferences to human intestine, thus promoting zoonosis.

The results of the present study could have implications toward the therapeutic applications targeting SadP and *S. suis* systemic infections (summarized in Fig 9). The gene encoding SadP type P_N_ has a downstream homolog, which is a pseudogene (has a stop codon). It could be speculated, that the gene encoding SadP type P_N_ is a putatively duplicated gene, which has evolved to bind Gb4 (Figure 9A). This study shows the molecular mechanism how N285 contributes to the binding of terminal GalNAcβ1-saccharide of Gb4 by a specific interaction with GalNAc hexose ring oxygen (Figure 9BC). The analogous N285-mediated binding mechanisms to Gb4 and glycomimetic phenylurea-galabiose could be utilized in the design of therapeutics against the invasive systemic infections without disturbing the commensal *S. suis* bacteria colonizing pig upper respiratory tract.

**Fig 9.**
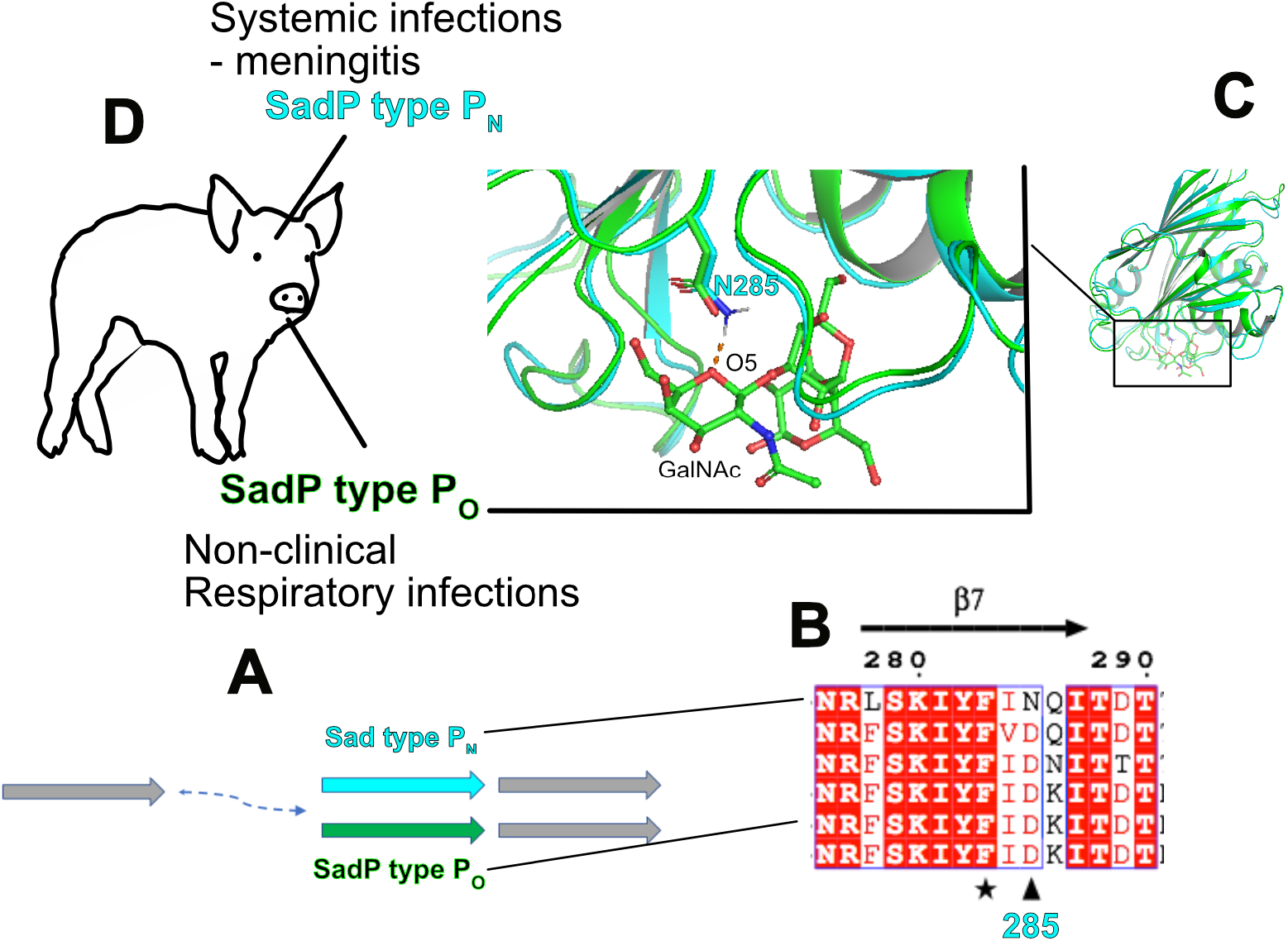
Summary of the results. **A.** SadP type P_N_ (cyan) and type P_O_ (green) are putative duplicates of the downstream SadP homolog (grey). **B.** Conservative change of type P_N_ SadP asparagine-285 to aspartic acid of SadP type P_O_. **C.** This study has shown the mechanism how N285 contributes to the binding of terminal GalNAcβ1-saccharide of Gb4 by a specific interaction with GalNAc hexose ring oxygen (orange dashed line). The 3D-structure of SadP type P_O_ from type 4 *S. suis* 6407 was obtained by homology modelling (Swiss Prot) and superimposed structures of type P_N_ (cyan) in complex with GalNAcβ1–Galα1–4Gal and type P_O_ (green) were created with Pymol. The structures suggest that the abolishment of Gb4 binding is based on the same mechanism as shown with N285D mutant. **D.** The results of the binding mechanisms of SadP type P_N_ with Gb4 and the distribution of SadP subtypes in *S. suis* clinical and non-clinical strains suggest that SadP Gb4 binding is associated with strains causing systemic disease. Therefore, the molecular mechanism of SadP type P_N_ binding to Gb4 could be targeted to prevent *S. suis* systemic diseases.

## Materials and Methods

### Bacterial strains

The chromosomal DNA was isolated from type P_N_ and P_O_ *S. suis* strains as described before (6) using GelElute bacterial genomic DNA kit according to the manufacturer’s instructions. *Streptococci* were grown in Todd Hewitt broth supplemented with 0.5 % yeast extract or in Columbia agar plates supplemented with 5 % sheep blood in 5 % CO_2_ at 37°C and *E. coli* strain were grown in LB medium. Antibiotics used were 30 μg/ml (pET28) and 100 μg/ml ampicillin pET46EkLIC) in *E. coli* and 500 μg/ml kanamycin for *S. suis* D282ΔSadP mutant (6).

*Escherichia coli* NovaBlue (Novagen) was used for the cloning of SadP constructs and strain BL21(DE3) for expression of the adhesin.

### Cloning and construction of recombinant adhesins

The galabiose binding *N*-terminal domains of SadP from hemagglutination subtypes P_N_ and P_O_ were cloned into pET46EkLIC vector as follows: The primer pairs SadPDER48 (gacgacgacaagatagaatcgctagaaccagatgtt) and SadPDER49 (gaggagaagcccggtttattcttctcaagggtaatctc) were designed to clone the 882 bp fragment of the adhesin *N*-terminal galabiose binding domain. The fragments were amplified with Phusion HotStart II DNA polymerase and were cloned into LIC-vector (LIC, ligation independent cloning) pET46EkLIC (Novagen). The ligation products were transformed into NovaBlue competent cells. This construct yielded a 33.4 kDa 6xHis-tagged fusion protein (SadP(31-328)). The sequence of *N*-terminal domains of SadP homologs were verified by sequencing with T7 promoter and T7 terminator primers. The vectors were transformed into expression strain BL21(DE3).

Site-specific mutant SadP(31-328) W258A was constructed by PCR with 5’-phosphorylated forward primer tgg aac gca tct gct ggt caa gct and reverse primer ttg aga att aaa acc att ttc. The PCR product was amplified with Phusion HotStart II DNA polymerase and the PCR fragment was ligated with T4 ligase (Promega) and transformed into Novablue competent cells. The resulting plasmid SadP(31-328)W258A-pET28a was sequenced to verify the mutation.

Synthetic gene construct and the corresponding site-specific mutants for WT SadP(125-328) type P_N_ were obtained from Genscript. Site-specific mutants constructed were Δ244-246 (PSAD-1), N285D (PSAD-2), E292Q (PSAD-3), N285D&Δ244-246 (PSAD-4) and H216-D246delinsDELFNRFPNKVDSTNNGDGAPFRFFNKE (PSAD-5) (replacement of aa 216-242 into corresponding sequence of type P_O_ of strain 6107).

### Bioinformatics

To compare the distribution of SadP alleles in *S. suis* from different infections e.g. invasive, respiratory and non-clinical samples, the presence of either P_N_ or P_O_ allele was detected from 374 previously published *S. suis* isolates (12) using SRST2 software version 0.2.0 with default settings (29). The SadP genes from *S. suis* P1/7 (AM946016.1) and 6407 (NZ_CP008921.1) were used as reference sequences for P_N_ and P_O_, respectively. The distribution of P_N_ and P_O_ alleles in the *S. suis* phylogeny was investigated by creating a core-genome maximum-likelihood tree using Roary version 3.12.0 (30) and RAxML version 8.2.8 (31) from the 374 isolates.

### Expression and purification of SadP proteins

The bacteria were grown at 30°C, 250 rpm, to an OD600 of 0.5, and the protein expression was induced with 0.2 mM IPTG for 3.5 hours. Bacteria were harvested by centrifugation with 3000 x g, at +4°C and stored at −84°C. The recombinant protein was purified with Ni-NTA affinity chromatography. Briefly, bacteria were lysed with 0.4 mg/ml hen egg lysozyme (Sigma) in 50 mM sodium phosphate buffer pH 8.0, containing 0.5 M NaCl, EDTA-free protease inhibitor cocktail (Pierce), 20 mM imidazole, 20 μg/ml deoxyribonuclease and 1 mM MgCl_2_ on ice for 30 mins. The lysate was sonicated to further homogenize the cell debris and was centrifuged at 20000 x g, +4°C for 30 min. The filtered lysate was purified with Ni-NTA affinity chromatography using HiPrep FF column connected to Äktaprime plus, GE Healthcare at +25°C. Further purification was done with a gel filtration HiLoad 16/60 Superdex 200 column using Tris-HCl, pH 7.5, 0.15 M NaCl as running buffer. The purity of the recombinant proteins was analyzed with SDS-PAGE chromatography. For native gel electrophoresis, the recombinant proteins were diluted to sample buffer without SDS and were separated with 8 % polyacrylamide gel.

### Crystallization

SadP N285D mutant (15 mg/ml) was crystallized with the hanging-drop vapor-diffusion method. The well condition contained 1.3-1.5 M sodium citrate tribasic, 0.1 M sodium cacodylate, pH 6.5. Equal volumes of well solution and protein (2 μL+2μL) were mixed and the crystals grew at 16 °C, initially as haystacks of very thin needles. Single crystals suitable for data collection were subsequently grown by using the seeding technique. A small part of the haystack was crushed with a metallic needle in the crystallization drop and 0.2 μL were diluted in 20 μL of mother liquor to create a seeding stock. Pre-equilibrated (1-d old) crystallization drops were seeded with 0.1 μL of the seeding stock. Single crystals started to grow after ~3 days.

### Data collection and crystal structure determination

Diffraction data were collected on the BioMAX beamline at MAX IV (Lund, Sweden) from a single crystal under cryogenic temperatures (100K). The crystal diffracted to 1.85 Å but the resolution was later truncated during processing to 2.05 Å owing to completeness. The crystal was found to belong to the *C*2 space group. Data processing was carried out with automated procedures in EDNA (32) and scaling was done with AIMLESS (33).

Molecular replacement with PHASER (34) was carried out to obtain initial phases using the native structure as a template. Five molecules in the crystallographic asymmetric unit were located (Matthews coefficient 2.67 Å^3^/Da corresponding to ~53.9% solvent content). Initial building of the structure was carried out using the automated building procedure in BUCCANEER (35) using the ccp4i2 interface (36). After the initial building that produced an almost complete structure, the refinement continued in PHENIX v.1.17.1-3660 (37) using simulated annealing at 1000K with maximum likelihood as target function. The first round of simulated annealing resulted in *R*_work_/*R*_free_ of 0.227/0.269. Inclusion of waters and rounds of manual rebuilding using COOT (38) alternated with PHENIX refinement resulted in the final structure with *R*_work_/*R*_free_ of 0.174/0.221 (Table S1). A glycerol molecule was found bound in each of the A, C, and D subunits. Two glycerol molecules were located in the binding site of subunit B. The structure has been deposited to the Protein Data Bank under the accession code 6YRO

### Isolation of SadP binding glycosphingolipids from porcine lung

Non-acid glycosphingolipids were isolated from porcine lung as described (39). Briefly, the lung tissue was lyophilized, and then extracted in two steps in a Soxhlet apparatus with chloroform and methanol (2:1 and 1:9, by volume, respectively). The material obtained was subjected to mild alkaline hydrolysis and dialysis, followed by separation on a silicic acid column. Acid and non-acid glycosphingolipid fractions were obtained by chromatography on a DEAE-cellulose column. In order to separate the non-acid glycolipids from alkali-stable phospholipids, this fraction was acetylated and separated on a second silicic acid column, followed by deacetylation and dialysis. Final purifications were done by chromatographies on DEAE-cellulose and silicic acid columns.

The total non-acid glycosphingolipid fraction (9 mg) from porcine lung was first separated on an Iatrobeads (latrobeads 6RS-8060; Iatron Laboratories, Tokyo) column (1.0 g) and eluted with increasing volumes of methanol in chloroform. Aliquots of the fractions obtained were analyzed by thin-layer chromatography. Fractions that were coloured green by anisaldehyde were tested for binding of SadP using the chromatogram binding assay. SadP bound to a fraction containing glycosphingolipids migrating in the tri-and tetraglycosylceramide regions on thin-layer chromatograms. This fraction (1.0 mg) was further separated on an Iatrobeads column (1.0 g), eluted with chloroform/methanol/water 65:25:4 (by volume), 30 x 0.5 ml, followed by chloroform/methanol/water 65:25:4, 10 mi. The SadP binding triglycosylceramide was eluted in fractions 12-17, and these fractions were pooled into one fraction migrating as a single band (0.2 mg, denoted fraction PL-1), and fraction migrating as a double band (0.3 mg, denoted fraction PL-<u>2</u>) on thin-layer chromatograms. In addition, a SadP tetraglycosylceramide was present in fractions 19-31, and pooling of these fractions gave 0.4 mg (denoted fraction PL-4).

### Reference glycosphingolipids

Total acid and non-acid glycosphingolipid fractions were isolated as described (39), and the individual glycosphingolipids were obtained by repeated chromatography on silicic acid columns, and by HPLC, and identified by mass spectrometry (20, 40) and 1H NMR spectroscopy (41).

### Thin-layer chromatography

Aluminum- or glass-backed silica gel 60 high performance thin-layer chromatography plates (Merck, Darmstadt, Germany) were used for thin-layer chromatography, and chromatographed with chloroform/methanol/water (60:35:8 by volume) as solvent system. The different glycosphingolipids were applied to the plates in quantities of 4 μg of pure glycosphingolipids, and 20-40 μg of glycosphingolipid mixtures. Chemical detection was done with anisaldehyde (42).

### Radiolabeling

Aliquots of 100 μg of the different SadP protein preparations were labeled with ^125^I by the Iodogen method according to the manufacturer’s instructions (Pierce/Thermo Scientific), giving approximately 2000 cpm/μg protein.

### Chromatogram binding assays

Binding of radiolabeled proteins to glycosphingolipids on thin-layer chromatograms was done as described (43). Chromatograms with separated glycosphingolipids were dipped for 1 min in diethylether/n-hexane (1:5, by volume) containing 0.5% (w/v) polyisobutylmethacrylate (Aldrich Chern. Comp. Inc., Milwaukee, WI). After drying, the chromatograms were soaked in PBS containing 2% (w/v) bovine serum albumin, 0.1% (w/v) NaN_3_ and 0.1% (w/v) Tween 20 (Solution A), for 2 h at room temperature. Thereafter the plates were incubated with ^125^I-labeled SadP protein (l-5 x 10^6^ cpm/ml) for 2 h at room temperature. After washing six times with PBS, and drying, the thin-layer plates were autoradiographed for 12 h using XAR-5 x-ray films (Eastman Kodak, Rochester, NY).

### Microtiter well assay

Binding of radiolabeled SadP to glycosphingolipids in microtiter wells was performed as described (43). In short, 250 μg of pure glycosphingolipids in methanol were applied to microtiter wells (Falcon 3911, Becton Dickinson Labware, Oxnard, CA). When the solvent had evaporated, the wells were blocked for 2 h at room temperature with 200 μl of BSA/PBS. Thereafter, the wells were incubated for 4 h at room temperature with 50 μl of ^125^I-labeled SadP (2 x 103 cpm/μl) diluted in BSA/PBS. After washing 6 times with PBS, the wells were cut out and the radioactivity was counted in a gamma counter.

### LC-ESI/MS of native glycosphingolipids

Native glycosphingolipids were analyzed by LC-ESI/MS as described (44). Glycosphingolipids were dissolved in methanol:acetonitrile in proportion 75:25 (by volume) and separated on a 200×0.150 mm column, packed in-house with 5 μM polyamine II particles (YMC Europe GmbH, Dinslaken, Germany). An autosampler, HTC-PAL (CTC Analytics AG, Zwingen, Switzerland) equipped with a cheminert valve (0.25 mm bore) and a 2 μl loop, was used for sample injection. An Agilent 1100 binary pump (Agilent technologies, Palo Alto, CA) delivered a flow of 250 μl/min, which was split down in an 1/16” microvolume-T (0.15 mm bore) (Vici AG International, Schenkon, Switzerland) by a 50 cm x 50 μm i.d. fused silica capillary before the injector of the autosampler, allowing approximately 2-3 μl/min through the column. Samples were eluted with an aqueous gradient (A:100% acetonitrile to B: 10 mM ammonium bicarbonate). The gradient (0-50% B) was eluted for 40 min, followed by a wash step with 100% B, and equilibration of the column for 20 min. The samples were analyzed in negative ion mode on a LTQ linear quadropole ion trap mass spectrometer (Thermo Electron, San José, CA), with an IonMax standard ESI source equipped with a stainless steel needle kept at – 3.5 kV. Compressed air was used as nebulizer gas. The heated capillary was kept at 270 °C, and the capillary voltage was –50 kV. Full scan (*m/z* 500-1800, two microscans, maximum 100 ms, target value of 30,000) was performed, followed by data-dependent MS^2^ scans (two microscans, maximun 100 ms, target value of 30.000) with normalized collision energy of 35%, isolation window of 2.5 units, activation q= 0.25 and activation time 30 ms). The threshold for MS^2^ was set to 500 counts. Data acquisition and processing were conducted with Xcalibur software version 2.0.7 (Thermo Fisher Scientific). Manual assignment of glycosphingolipid sequences was done with the assistance of the Glycoworkbench tool (Version 2.1), and by comparison of retention times and MS^2^ spectra of reference glycosphingolipids.

### Endoglycoceramidase digestion and LC-ESI/MS

Endoglycoceramidase II from *Rhodococcus* spp. <u>(Ito)</u> (Takara Bio Europe S.A., Gennevilliers, France) was used for hydrolysis of glycosphingolipids. Briefly, 50 μg of glycosphingolipids were resuspended in 100 μl 0.05 M sodium acetate buffer, pH 5.0, containing 120 μg sodium cholate, and sonicated briefly. Thereafter, 1 mU of enzyme was added, and the mixture was incubated at 37°C for 48 h. The reaction was stopped by addition of chloroform/methanol/water to the final proportions 8:4:3 (by volume). The oligosaccharide-containing upper phase thus obtained was separated from detergent on a Sep-Pak QMA cartridge (Waters). The eluant containing the oligosaccharides was dried under nitrogen and under vacuum.

The glycosphingolipid-derived oligosaccharides were resuspended in 50 μl of water and analyzed by LC-ESI/MS as described (20). The oligosaccharides were separated on a column (200 x 0.180 mm) packed in-house with 5 μm porous graphite particles (Hypercarb, Thermo-Hypersil, Runcorn, UK). An autosampler, HTC-PAL (CTC Analytics AG, Zwingen, Switzerland) equipped with a cheminert valve (0.25 mm bore) and a 2 μl loop, was used for sample injection. An Agilent 1100 binary pump (Agilent technologies, Palo Alto, CA) delivered a flow of 250 μl/min, which was split down in an 1/16” microvolume-T (0.15 mm bore) (Vici AG International, Schenkon, Switzerland) by a 50 cm x 50 μm i.d. fused silica capillary before the injector of the autosampler, allowing approximately 2-3 μl/min through the column. The oligosaccharides (3 μl) were injected on to the column and eluted with an acetonitrile gradient (A: 10 mM ammonium bicarbonate; B: 10 mM ammonium bicarbonate in 80% acetonitrile). The gradient (0-45% B) was eluted for 46 min, followed by a wash step with 100% B, and equilibration of the column for 24 min. A 30 cm x 50 μm i.d. fused silica capillary was used as transfer line to the ion source.

The oligosaccharides were analyzed in negative ion mode on an LTQ linear quadrupole ion trap mass spectrometer (Thermo Electron, San José, CA). The IonMax standard ESI source on the LTQ mass spectrometer was equipped with a stainless-steel needle kept at –3.5 kV. Compressed air was used as nebulizer gas. The heated capillary was kept at 270 °C, and the capillary voltage was –50 kV. Full-scan (*m/z* 380-2 000, 2 microscans, maximum 100 ms, target value of 30 000) was performed, followed by data dependent MS^2^ scans of the three most abundant ions in each scan (2 microscans, maximum 100 ms, target value of 10 000). The threshold for MS^2^ was set to 500 counts. Normalized collision energy was 35%, and an isolation window of 3 u, an activation q = 0.25, and an activation time of 30 ms, was used. Data acquisition and processing were conducted with Xcalibur software (Version 2.0.7).

Manual assignment of glycan sequences was done on the basis of knowledge of mammalian biosynthetic pathways, with the assistance of the Glycoworkbench tool (Version 2.1), and by comparison of retention times and MS^2^ spectra of oligosaccharides from reference glycosphingolipids (20).

### S. suis cell binding assay

EA.hy926 cells were maintained and grown in DMEM, 10 % FCS medium at 37°C, 5 % CO_2_. For bacterial binding assay the cells were detached with Trypsin-EDTA, counted and 15 000 cells/well added into round glass coverslips in 24 well-plates. Cells were grown for 24 - 48 h to subconfluency.

*S. suis* D282 WT strain and D282-ΔsadP were grown in THY overnight at 37°C, 5 % CO_2_. Bacteria were diluted 1/20 into prewarmed 37°C THY and were grown to OD550 of 0.2 and were diluted 1/100 into prewarmed DMEM, 10 % FCS without antibiotics. 500 μl of bacterial dilution was pipetted into the wells (MOI 100:1) and the plate was centrifuged with 800 x g, 15 min at 20°C.

The plate was incubated at 37°C for 1 h. The wells were washed 4 x 1000 μl of PBS. The cells were stained with DiffQuick kit, washed and the coverslips were mounted on glass slides with Permount.

The bacterial binding was quantitated by microscopy with 100 x objective with immersion oil. The bound bacteria were enumerated by counting two wells for each sample. The results were expressed as average of bacteria / optical field.

For flow cytometry, N-terminal domain of SadP was labelled with FITC (Sigma-Aldrich) in 0.2 M borate buffer, pH 8.0, 0.16 M NaCl (50:1 molar ratio of FITC/SadP, 83 nmol of SadP and 1.7 μmol of FITC) and the labelling was stopped with Tris-buffer. The labelled SadP was purified from the free FITC with PD-10 (GE Healthcare) desalting column. The EA.hy926 cells were seeded into 6 well culture dishes (1 x 10^5^ cells/well) and were grown for 48 h. The cells were washed with DMEM and were incubated with 400 ng/ml of labelled SadP with or without 10 μg/ml pigeon ovomucoid. After binding, the wells were washed with phosphate Buffered Saline (PBS, 0.15 M NaCl, 2.7 mM KCl, 8.1 mM Na_2_HPO_4_, 1.5 mM KH_2_PO_4_). and were fixed with 2 % (w/v) paraformaldehyde in PBS for 15 min. The cells were washed twice with PBS and the cells were scraped of the wells and suspended into the PBS. 10 000 cells were analyzed for SadP binding with FACSCalibur.

### Amplified luminescent proximity homogeneous assay (AlphaScreen)

The AlphaScreen assay was optimized as described before (19). Briefly, the assay was performed using AlphaScreen Streptavidin Donor beads and NiNTA Acceptor beads (PerkinElmer). The molar concentrations of galabiose-containing biotinylated ovomucoid (receptor) and adhesin were optimized by finding the hook point for the interaction by setting up a matrix of proteins in 96 well AlphaPlates (Perkin Elmer) in 10 mM Tris-HCl, pH 7.5, 0.15 M NaCl, 0.2% BSA, 0.05 % Tween 20 0.2% (TBST-0.2%BSA). For the optimized assay dilutions of 5 μl of the His-tagged adhesin and 5 μl of biotinylated ovomucoid were pipetted into the wells and the plates were centrifuged for 1 min, 1000 x g. The plates were incubated at +25°C for 1 hr. This was followed by addition of Ni-NTA-beads (20 μg/ml) and incubation for 1.5 hrs. Finally, streptavidin donor beads were added (20 μg/ml) and the mixtures were incubated for 30 mins and were then measured with the Ensight multimode reader (Perkin Elmer) using excitation wavelength of 680 nm with donor beads and measurement of emission wavelength of 615 nm from AlphaLISA anti-HIS acceptor beads. For inhibition assays the oligosaccharides, whose structures have been described before (25) or depicted in Figure 4 for TMSEt-glycosides of GalNAcβ1–3Galα1–4Gal and 3Galα1–4Gal, were diluted into TBST-0.2%BSA buffer and were mixed with the His-tagged adhesin and biotinylated ovomucoid. The binding was measured as described above. The binding inhibition data was fitted using Prism with settings of log(inhibitor) vs. response slope (four parameters).

### Isothermal titration calorimetry

SadP-D282(31-328) (P_N_), SadP-6107(31-328) (P_O_), SadP(125-329) type P_N_ and site-specific mutant SadP(125-328)N285D were desalted into PBS using PD-10 desalting column (GE Healtcare). The synthetic oligosaccharides were diluted to the same buffer. 0.1 - 0.2 μM of SadP proteins in a 350 μl vessel was titrated with 1.5 - 2 mM solutions of galabiose oligosaccharides using MicroCal (Malvern). The protein solution was stirred with 750 rpm at 25°C and 16 injections of a volume of 2.49 μl were injected at 180 s intervals. The data from single determinations was analyzed with Origin.

### Preparation of liposomes and SadP binding assay

Vesicles were prepared as described before (45). Briefly, POPC (1-palmitoyl-2-oleoyl-glycero-3-phosphocholine, Avanti Polar lipids), cholesterol and glycolipids were dissolved and mixed into the chloroform : methanol (2:1, vol/vol) to contain 30 % cholesterol and 2 mM glycolipids. The solvent was evaporated and the lipid film was suspended for 30 min at 60 °C in 10 mM Tris-Cl, 140 mM NaCl buffer. The lipid-buffer suspension was briefly vortexed followed by the extrusion procedure (Avanti mini extruder using 0.1 μm polycarbonate membranes filter, (Avanti Polar Lipids, Alabaster, AL, USA)) to form large unilamellar vesicles.

PVDF membrane was wetted (wetting in 100 % MeOH, washing 4 x 5 min MilliQ water) and was overlaid onto wet Whatman filter paper. Liposomes were serially diluted 1/10 into Tris-buffered saline and 2 μl of liposomes containing 2 mM, 0.2 mM and 0.02 mM of glycolipid was pipetted onto the membrane and the membrane was not allowed to dry during the pipetting. The membrane was saturated with 2 % BSA (w/v) in TBS for 1 h at 20°C and was incubated with 10 μg/ml recombinant SadP for 2 h at 20°C. The membrane was washed 4 x 5 min with TBS and the bound SadP was detected with 1:10000 dilution of anti-His primary antibody (Sigma) and 1:20000 dilution of HRP-labelled secondary rabbit anti-mouse antibody (DakoCytomation). The membrane was incubated with ECL substrate (WesternBright Quantum, Labtech) and the SadP binding was imaged with Fuji LAS-4000.

### STD-NMR

STD-NMR measurements were carried out on a Bruker AVANCE III HD spectrometer (Bruker BioSpin GmbH, Rheinstetten, Germany) operating at 600.13 MHz, equipped with a nitrogen-cooled triple-channel TCI inverse CryoProbe™. The pulse sequence used for the experiments was the stddiffesgp provided by Bruker. A 1x PBS buffer with a pH of 7.45 with 10% D_2_O, containing 50 μM protein and 2500 μM Gb3 trisaccharide was used for the experiments. The on-resonance pulse was set to the methyl groups at 0.9 ppm and the off-resonance pulse was set at 20 ppm. A saturation time of 2 s was used and the experiments were carried out at 313 K to somewhat improve the weak signals, likely resulting from slow k_off_. A blank experiment with only Gb3 and no protein was done to rule out any self-STD.

### Molecular dynamics simulations

Molecular dynamics simulations were performed with the OPLS3 force field in Desmond (Schrödinger Release 2019-4: Desmond Molecular Dynamics System, D. E. Shaw Research, New York, NY, 2017; Maestro-Desmond Interoperability Tools, Schrödinger, New York, NY, 2017) using default settings except for the length of the simulation. Simulations were performed with periodic boundary conditions using orthorhombic simulation boxes with SPC water model. Counter ions were used to neutralize each other. Starting conformations of ligands in complex with SadP (pdb id 5BOA) were built by manually placing ligand galabiose residues in an identical position to that in the crystal structure (pdb id 5BOA). Complexes with the mutants were generated by exchanging N285 for D285. The complexes were then subjected to 100 ns molecular dynamics simulations. The complex of the phenylurea derivative with the N285D mutant was not stable during the 100 ns simulations. Therefore, light constraints (1 kcal·mol^−1^·Å^−1^) on the protein secondary structures were applied. Molecular images were generated using PyMOL v2.30 (Schrodinger LLC).

## Author contributions

MMJ and ST purified, characterized the glycolipids and performed TLC overlay experiments. TK and JC performed the bioinformatic analysis. EB and SH purified recombinant WT and site-specifically mutated SadP proteins, optimized and performed AlphaScreen analysis and competitive inhibition assays. SH performed the cell binding and flow cytometry analysis. SM and TN prepared Gb3 and Gb4 containing liposomes. ACP performed the structural analysis and modelling of SadP. APS and UJN did the molecular dynamics calculations. JR and RL performed the STD-NMR experiments. JK and SH did the ITC experiments. JF, ST and SH planned the project. APS, UJN, JF and JC contributed to the editing of the article. ST and SH wrote the manuscript.

## Acknowledgements

The synthetic glycolipid Galα1–4Galβ1–O-bis-(SO_2_-C_16_H_33_)_2_ was a kind gift from Professor Göran Magnusson, Lund University, Sweden. ACP thanks Noushin Madani for help during crystallographic data collection. Infrastructure support from Biocenter Finland is acknowledged. SH thanks Aki Stubb for purification of recombinant SadP proteins. Infrastructure support from Biocenter Finland is acknowledged. Access to MAX IV was financially assisted by iNEXT, project number 653706, funded by the Horizon 2020 programme of the European Union. TK and JK were funded by ERC grant no. 742158. SH and JF were funded by project funding from the Magnus Ehrnrooth Foundation, Turku University Foundation, ST from the Swedish Cancer Foundation (ST; No. 18 0760), and by governmental grants to the Sahlgrenska University Hospital.

## Supporting information

**S1 Fig. LC-ESI/MS of the native fraction PL-1 from pig lung.** (A) Base peak chromatogram from LC-ESI/MS of fraction PL-2 from pig lung. (B) MS^2^ of the ion at *m/z* 1132 (retention time 14.1 min). The interpretation formula shows the deduced glycosphingolipid structure.

**S2 Fig. LC-ESI/MS of the native fraction PL-2 from pig lung. (A) Base** peak chromatogram from LC-ESI/MS of fraction PL-2 from pig lung. (B) Mass chromatogram of *m/z* 1022. (C) Mass chromatogram of *m/z* 1132. (D) MS^2^ of the ion at *m/z* 1022 (retention time 13.1 min). (E) MS^2^ of the ion at *m/z* 1132 (retention time 12.5 min). (F) Interpretation formulas showing the deduced glycosphingolipid structures.

**S3 Fig. LC-ESI/MS of the oligosaccharides obtained by digestion of fraction PL-1, PL-2, reference globotriaosylceramide, and reference isoglobotriaosylceramide with *Rhodococcus* endoglycoceramidase II.** A. Base peak chromatogram from LC-ESI/MS of the oligosaccharides derived from fraction PL-1. B. Base peak chromatogram from LC-ESI/MS of the oligosaccharides derived from fraction PL-2. C. Base peak chromatogram from LC-ESI/MS of the oligosaccharides derived from reference globotriaosylceramide (Galα1–4Galβ1–4Glcβ1–O–Cer). D. Base peak chromatogram from LC-ESI/MS of the oligosaccharides derived from reference isoglobotriaosylceramide (Galα1–3Galβ1–4Glcβ1–O–Cer). E. MS^2^ of the ion at *m/z* 503 (retention time 18.4 min) in (A). F. MS^2^ of the ion at *m/z* 503 (retention time 17.7 min) in (B). G. MS^2^ of the ion at *m/z* 503 (retention time 17.4 min) in (C). H. MS^2^ of the ion at *m/z* 503 (retention time 20.2 min) in (D). The interpretation formula shows the deduced oligosaccharide structure. The identification of oligosaccharides was based on their retention times, determined molecular masses and subsequent MS^2^ sequencing.

**S4 Fig. LC-ESI/MS of the oligosaccharides obtained by digestion of fraction PL-4, and reference globotetraosylceramide with *Rhodococcus* endoglycoceramidase II.** A. Base peak chromatogram from LC-ESI/MS of the oligosaccharides derived from fraction PL-4. B. MS^2^ of the ion at *m/z* 706 (retention time 18.4 min) in (A). C. MS^2^ of the ion at *m/z* 706 (retention time 18.6 min) from LC-ESI/MS of the oligosaccharides derived from reference globotetraosylceramide (GalNAcβ1–3Galα1–4Galβ1–4Glcβ1–O–Cer). D. MS^2^ of the ion at *m/z* 998 (retention time 19.4 min) in (A). E. MS^2^ of the ion at *m/z* 998 (retention time 21.4 min) in (A). The interpretation formula shows the deduced oligosaccharide structure. The identification of oligosaccharides was based on their retention times, determined molecular masses and subsequent MS^2^ sequencing.

**S5 Fig. SadP types P_N_ and P_O_ glycolipid binding specificity analysed with TLC overlay analysis and microwell binding assay: A**. Decreasing glycolipid dilutions from 2 to 0.2 μg per lane probed with SadP type P_N_ full length and it’s N-terminal domain SadP(31-328). **B.** Microwell binding assay with radiolabelled SadP type P_N_ to 250 ng of glycolipids, Gb2 galabiosylceramide, Gb2-S Galα1–4Galβ1–O-*bis*-(SO_2_-C_16_H_33_)_2_, Gb3 non-OH globotriaosylceramide with non-hydroxy ceramide, Gb3 OH globotriaosylceramide with hydroxy ceramide, iGb3 isoglobotriaosylceramide, Gb4, globotetraosylceramide, P1 pentaosylceramide. <u>Leb Lewis bhexaosylceramide</u>.

**S6 Fig.** SDS-PAGE of the type P_N_ WT SadP(125-329) and the site-specific mutants N285D and E292Q, deletion Δ244-246 and N285D&Δ244-246 mutants were constructed into the type P_N_ SadP(125-328) cloned into the plasmid pET28a. In addition, the 28 amino acid long region H216-D246 of the type P_N_ was replaced by the corresponding region of type P_O_.

**S7 Fig. A.** Binding of SadP wild type and site-specific mutants to Galα1–4Gal analysed with AlphaScreen **. B - F.** Inhibition assay for the binding of site-specific mutants of PSAD1-5 (PSAD-1 Δ244-246, PSAD-2 N285D, PSAD-3 E292Q, PSAD-4 N285D#x0026;Δ244-246 and PSAD-5 H216-D246delinsDELFNRFPNKVDSTNNGDGAPFRFFNKE (replacement of aa 216-242 into corresponding sequence of type P_O_ of strain 6107).

**S8 Fig. Structure analysis of TMSEt** (2-trimethylsilylethyl) glycosides

**S9 movie. MD simulation of type P_N_ WT SadP(125-329) and Galα1–4Gal**

**S10 movie. MD simulation of N285D SadP(125-329) and Gal**α**1–4Gal**

**S11 movie. MD simulation of of type P_N_ WT SadP(125-329) and GalNAcβ 1–3Galα1–4Gal**

**S12 movie. MD simulation of of type P_N_ WT SadP(125-329) and 3’-phenylurea-Galα1–4Gal**

**S13 movie. MD simulation of N285D SadP(125-329) and GalNAcβ1–3Galα1–4Gal**

**S14 movie. MD simulation of N285D SadP(125-329) and 3’-phenylurea-Galα1–4Gal**

