## Supplementary figures and images for "The binding mechanism of *Streptococcus suis* accessory virulence factor and adhesin SadP to globotetraosylceramide"

### S1 Fig

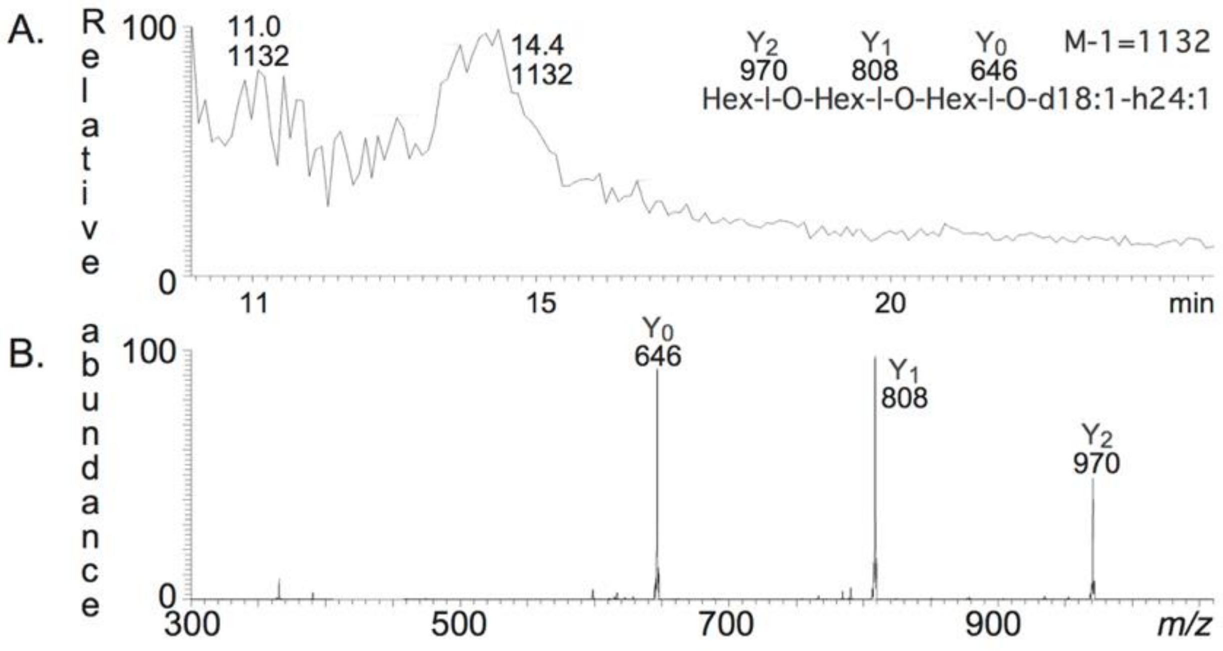

### S2 Fig

S2 Fig

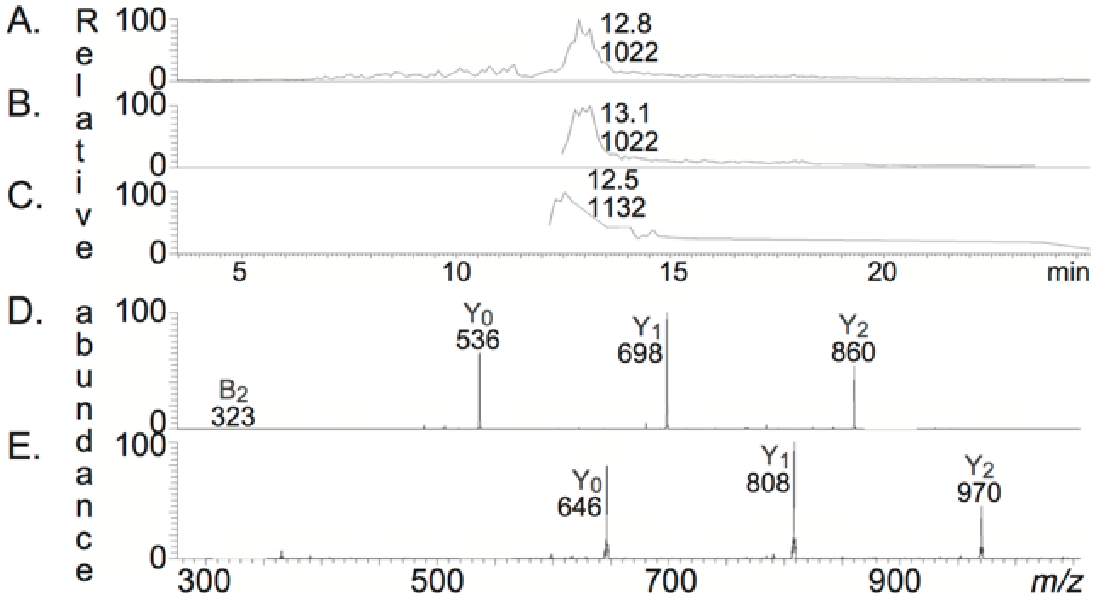


F.

$Y_2$ 860 — $Y_1$ 698 — $Y_0$ 536 — M-1=1022

Hex-l-O-Hex-l-O-Hex-l-O-d18:1-16:0

$Y_2$ 970 — $Y_1$ 808 — $Y_0$ 646 — M-1=1132

Hex-l-O-Hex-l-O-Hex-l-O-d18:1-24:1

### S3 Fig

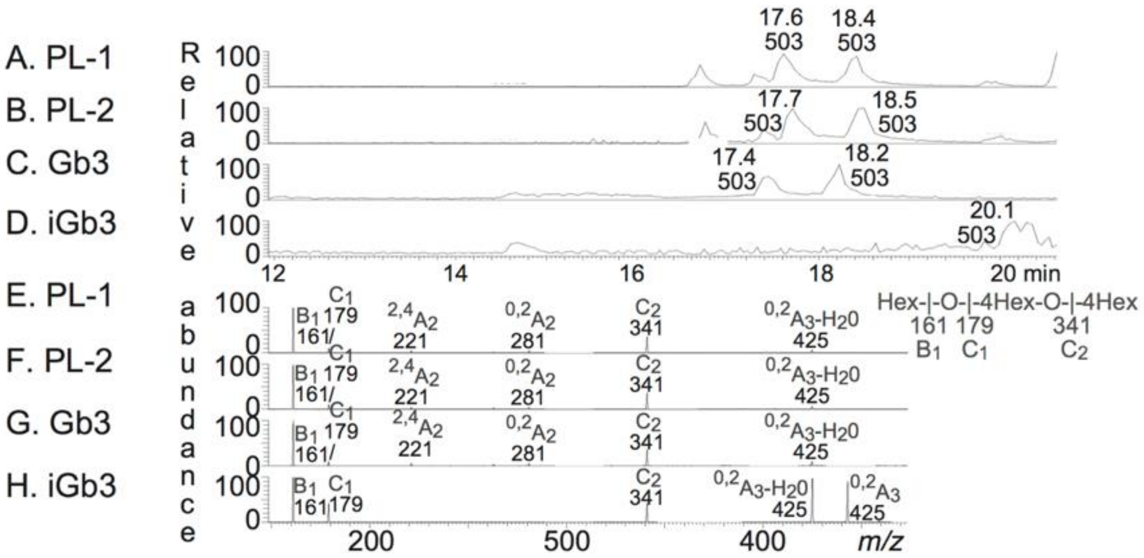

### S4 Fig

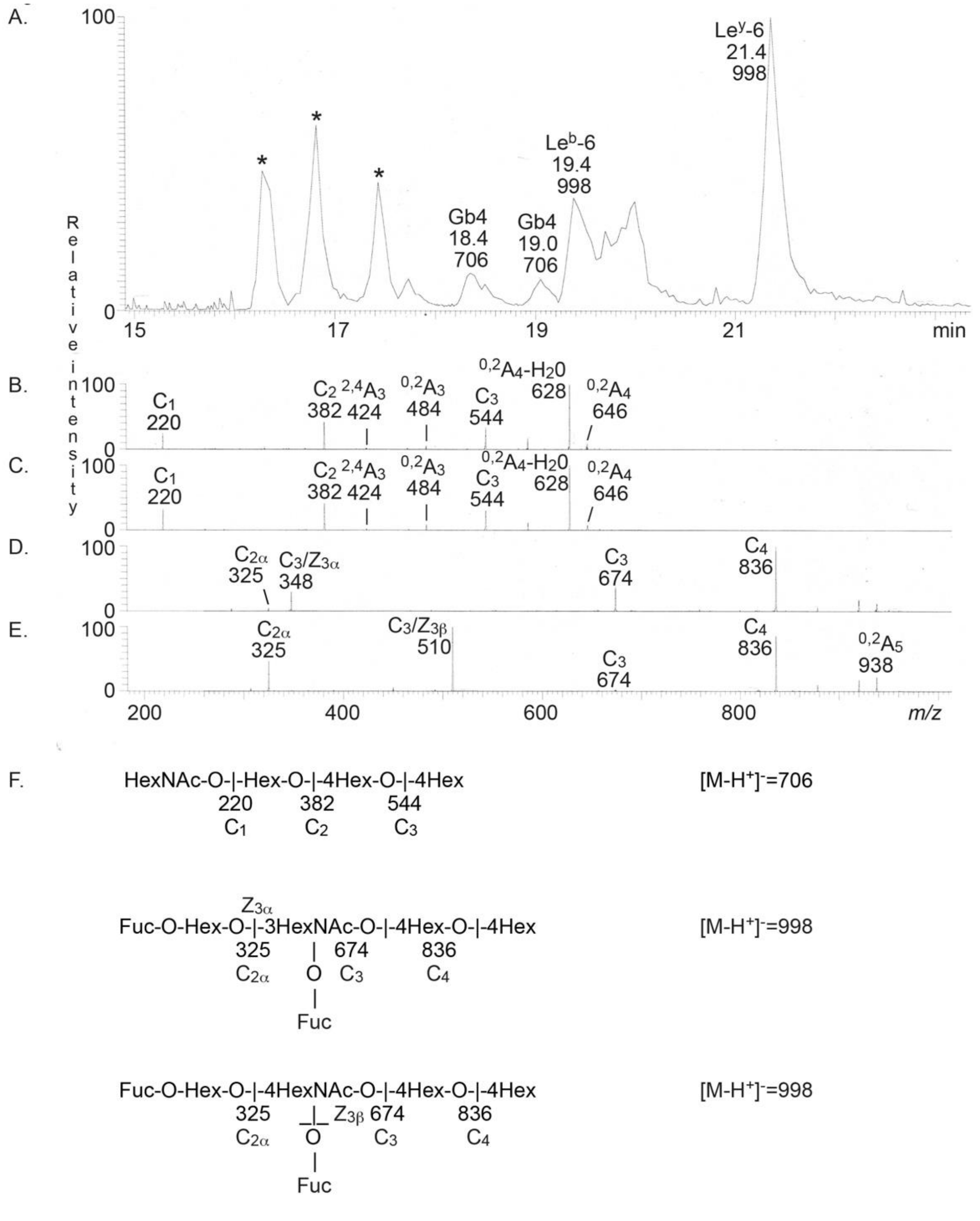

### S5 Fig

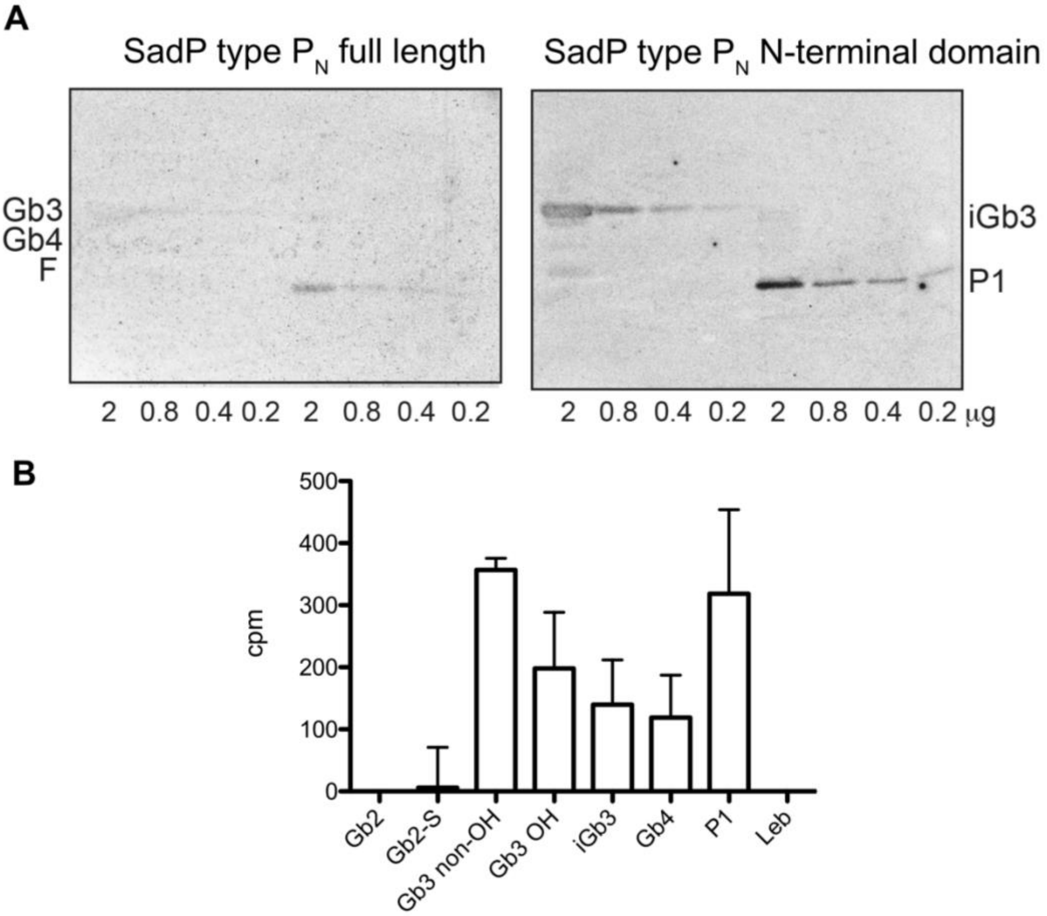

### S6 Fig

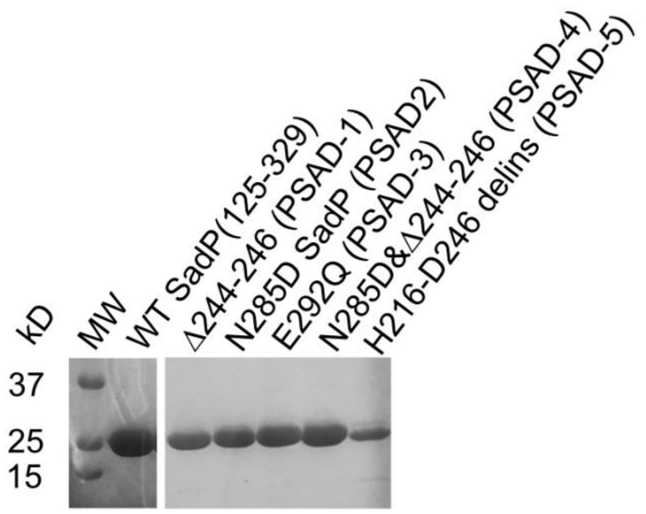

### S7 Fig

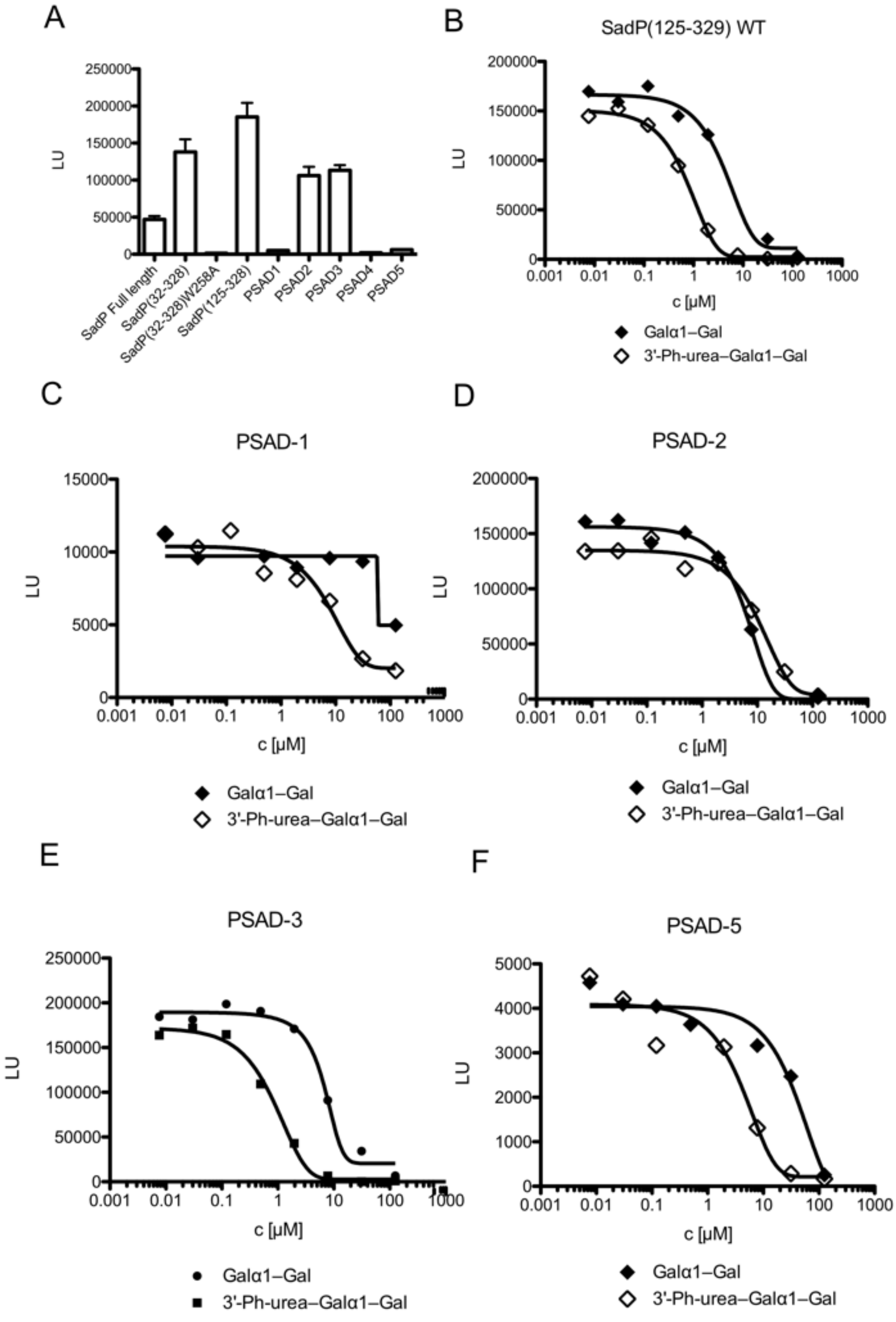

### S8 Fig

S8 Fig

Galα1–4Gal-O-TMSEt

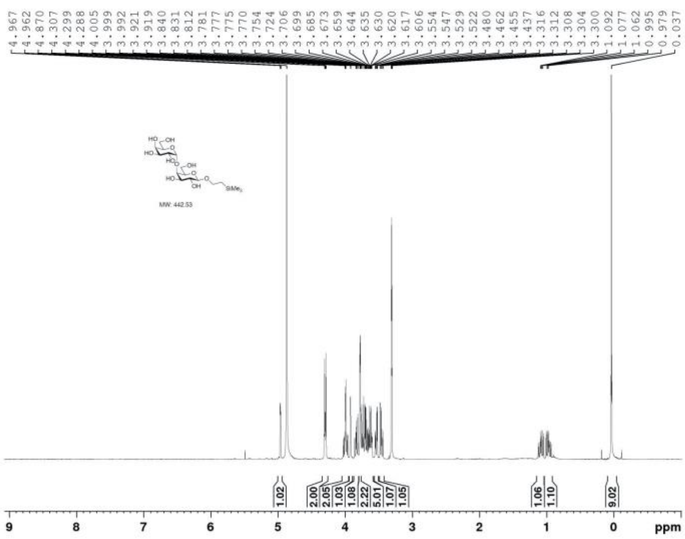

GalNAcβ1–3Galα1–4Gal-O- TMSEt

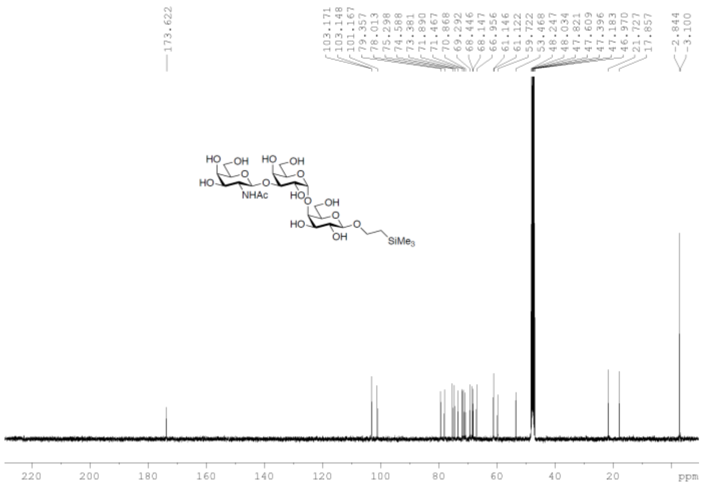
